# Amoeba predation of *Cryptococcus neoformans* results in pleiotropic changes to traits associated with virulence

**DOI:** 10.1101/2020.08.07.241190

**Authors:** Man Shun Fu, Livia C. Liporagi-Lopes, Samuel R. dos Santos Júnior, Jennifer L. Tenor, John R. Perfect, Christina A. Cuomo, Arturo Casadevall

## Abstract

Phagocytic amoeboid predators such as amoeba have been proposed to select for survival traits in soil microbes such as *Cryptococcus neoformans* that can also function in animal virulence by defeating phagocytic immune cells, such as macrophages. Several prior studies have shown that incubation of various fungal species with amoeba can enhance their virulence. However, the mechanisms by which fungi adapt to amoeba and thus change their virulence are unknown. In this study we exposed three strains of *C. neoformans* (1 clinical and 2 environmental) to predation by *Acanthamoeba castellanii* for prolonged periods of time and then analyzed surviving colonies phenotypically and genetically. Surviving colonies were comprised of cells that expressed either pseudohyphal or yeast phenotypes, which demonstrated variable expression of such traits associated with virulence such as capsule size, urease production and melanization. Phenotypic changes were associated with aneuploidy and DNA sequence mutations in some amoeba-passaged isolates, but not in others. Mutations in the gene encoding for the oligopeptide transporter (CNAG_03013; *OPT1*) were observed among amoeba-passaged isolates from each of the three strains. In addition, isolates derived from environmental strains gained the capacity for enhanced macrophage toxicity after amoeba selection and carried mutations on the CNAG_00570 gene, which encodes Pkr1 (AMP-dependent protein kinase regulator) but were less virulence in mice because they elicited more effective fungal-clearing immune responses. Our results indicate that *C. neoformans* survival under constant amoeba predation involves the generation of strains expressing pleiotropic phenotypic and genetic changes, which confer increase resistance against protozoal predation. Given the myriad of potential predators in soils the diversity observed among amoeba-selected strains suggests a bet-hedging strategy whereby variant diversity increases the likelihood that some will survive predation.

**Author summary:** *Cryptococcus neoformans* is a ubiquitous environmental fungus that is also a leading cause of fatal fungal infection in humans, especially among immunocompromised patients. Cryptococcosis is a worldwide concern due to its high mortality rate. A major question in the field is how an environmental yeast such as *C. neoformans* becomes a human pathogen when it has no need for animal host in its life cycle. Previous studies showed evidence that *C. neoformans* increases its pathogenicity after interacting with its environmental predator amoebae. Amoebae behave like macrophages, an important immune cell in human body, so it is considered as a training ground for pathogens to resist macrophages. However, how *C. neoformans* changes its virulence through interacting with amoebae is unknown. Here, we exposed *C. neoformans* to amoebae for a long period of time. We found that *C. neoformans* cells recovered from amoebae manifested numerous changes to phenotypes related to its virulence and one of the amoeba-passaged *C. neoformans* cells had enhanced ability to kill macrophages. We further analyzed their genome sequences and found various mutations in different cells of amoeba-passaged *C. neoformans*, showing that DNA mutations may be the major cause of the phenotypic changes after interacting with amoebae. Our study indicates that fungal survival in the face of amoeba predation is associated with the emergence of pleiotropic phenotypic and genomic changes that increase the chance of fungal survival.

## Introduction

*C. neoformans* is a major life-threatening fungal pathogen that predominantly infects severely immunocompromised patients and causes over 180,000 deaths per year worldwide [1]. C. *neoformans* expresses virulence factors that promote its pathogenicity in humans, including formation and enlargement of a polysaccharide capsule that interferes with the host immune system in varied ways, melanin production that protects against oxidative stress [2–5], and extracellular secretion of various enzymes including phospholipase and urease [6,7]. *C. neoformans* is found primarily and ubiquitously in environments such as soils contaminated with bird excreta or from trees [8–11]. It is a saprophyte and does not require an animal host for survival and reproduction. Besides, there is rare evidence of human-to-human transmission and thus it is unlikely that its virulence traits were selected for causing disease in humans or animals. That raises the fundamental question of how *C. neoformans* acquired those traits, which are essential for pathogenesis of cryptococcosis in human.

A hypothesis to solve this enigma is that of coincident selection, resulting from selective pressures in both natural environmental and animal niches such as predatory amoeba and nematodes [12]. According to this view, microbial traits selected for environmental survival also confer the capacity for virulence by promoting survival in animal hosts [12]. For example, the capsule can protect the fungi from desiccation and against predatory amoeba [13,14] while melanin may reduce damage of fungi from the exposure to UV radiation, osmotic stresses or extreme temperatures [15–18]. Urease provides a nutritional role involved in nitrogen acquisition in the environment [19]. Moreover, it is striking that *C. neoformans* isolates from the soil are virulent for animal hosts [20]. Understanding the evolutionary adaption of *C. neoformans* in nature will help us to understand further the origin of virulence and pathogenesis of cryptococcosis.

Amoebae are one of the major sources of selective pressure in nature for broad range of soil microorganisms that have pathogenic potential for humans, including bacteria such as *Legionella pneumophila, Mycobacterium spp* and fungi such as *Cryptococcus neoformans, Aspergillus fumigatus* and *Paracoccidioides spp* [14,21–24]. Similar to human macrophages, amoebae ingest microorganisms, undergo a respiratory burst, phagosome maturation and acidification, expresses cell surface receptors and expel undigested materials [25–31]. However, many bacteria and fungi have strategies to survive in amoebae, that function in parallel for survival in mammalian phagocytic cells. For example, *L. pneumophila* utilizes similar cellular and molecular mechanisms of invasion, survival and replication inside both amoeba and macrophages [32–37]. Amoeba-grown *L. pneumophila* are more invasive for epithelial cells and macrophages [21]. After passage in amoeba, *Mycobacterium avium* enhances both entry and intracellular replication in epithelial cells and is more virulent in the macrophage and mouse models of infection [22]. Among fungal pathogens, concordance of virulence factor function for amoeba and animals was also demonstrated for *A. fumigatus* [23]. For example, the mycotoxin fumagillin can inhibit the growth of *Entamoeba histolytica* while it can also cause mammalian epithelial cell damage [38]. Many studies have been done to explore amoeba-*C. neoformans* interaction, and shown evidence that amoebae influence the virulence of *C. neoformans* for mammalian infection [39,40].

*Acanthamoeba castellanii* was originally isolated from cultures of a *Cryptococcus* spp., and like other amoebae species preys on *Cryptococcus* spp [41,42]. There is evidence that amoebae are natural predators of *C. neoformans* in the natural environment [43]. On the other hand, *C. neoformans* is able to resist the destruction by amoeba, especially in nutrient poor conditions [44] without metal cations [45]. Several studies have shown that the virulence factors and the cellular process that fungi use for defending against amoeba predation are remarkably similar to those employed for mammalian virulence. For example, the capsule formation and melanin production are important for *C. neoformans* to resist predation by *A. castellanii* and play important roles for pathogenicity in mammalian infection [14,39]. Interestingly, the phospholipids that are secreted by both macrophages and amoebae trigger capsule enlargement [40]. The non-lytic exocytosis process which is found in macrophage containing *C. neoformans* can be also observed in *A. castellanii* and *D. discoideum* through the similar action of actin polymerization [46,47]. Transcriptional studies showed a conserved metabolic response of *C. neoformans* to the microenvironments of both macrophage and amoebae [48]. All those common strategies found to adapt to both amoebae and macrophages support the hypothesis that cryptococcal pathogenesis is derived from the interaction with amoebae in natural environment. More direct evidence comes from the experiment on the passage of an attenuated cryptococcal strain to *D. discoideum* cultures that shows enhancement of fungal virulence in a murine infection model [39]. Passaged *C. neoformans* also exhibits capsule enlargement and rapid melanization, suggesting that those are mechanisms to enhance the survival of fungus in mice. However, the underlying mechanism on how these phenotypic changes occur is still unclear. In this study, we sought to determine the long-term evolutionary adaption of *C. neoformans* when interacting with amoeba and whether the adaption affected virulence traits for animal hosts. Our results show that persistent amoeba predation was associated with the emergence of pleiotropic phenotypic changes of *C. neoformans*.

## Result

### Selection of amoeba-resistance strains

We studied the interaction between *C. neoformans* and *Acanthamoeba castellanii* by culturing them together on Sabouraud agar. For the initial experiments, we used the well-studied common laboratory strain H99. The experimental setup involved spreading approximately 200 cryptococcal cells on agar followed by placing approximately 5000 *A. castellanii* cells on the plate. After approximately month of co-incubation, small colonies emerged within the predator zone of *A. castellanii* (Fig 1A), sometimes under the mat of amoeba. Microscopic morphological analysis of cells in those colonies revealed pseudohyphal and hyphal forms of *C. neoformans* (Fig 1B & C). We selected 20 single hyphal cells from two colonies (ten hyphae from each colony) and these were transferred to a fresh Sabouraud agar plate without amoeba (Fig 1D, E & I). After 24 h, microcolonies composed exclusively of yeast cells emerged on the agar (Fig 1F & J), which manifested two distinct colonies morphologies, smooth and serrated, after two days of agar growth (Fig 1G & K). All of the cells from these colonies were yeasts (Fig 1H & L). The same experiment was then repeated with two environmental avirulent *C. neoformans* strains, A1-35-8 and Ftc555-1, but this time total 20 single hyphae were picked from four survival colonies (five hyphae from each colony) to a fresh agar plate. Like the experience with H99, these strains responded to the presence of amoeba by generating cells that formed colonies with various cellular and colony morphologies, of which some (A4-6) were slightly serrated with pseudohyphal cells (Fig 1M). We also observed some hyphal colonies formed by Ftc555-1 cells but eventually they converted back to yeast cells when streaked on fresh agar medium (Fig 1N). The results showed that after interacting with amoebae, *C. neoformans* can develop high variety of cellular and colony morphologies even in amoebae-free medium.

**Fig 1.**
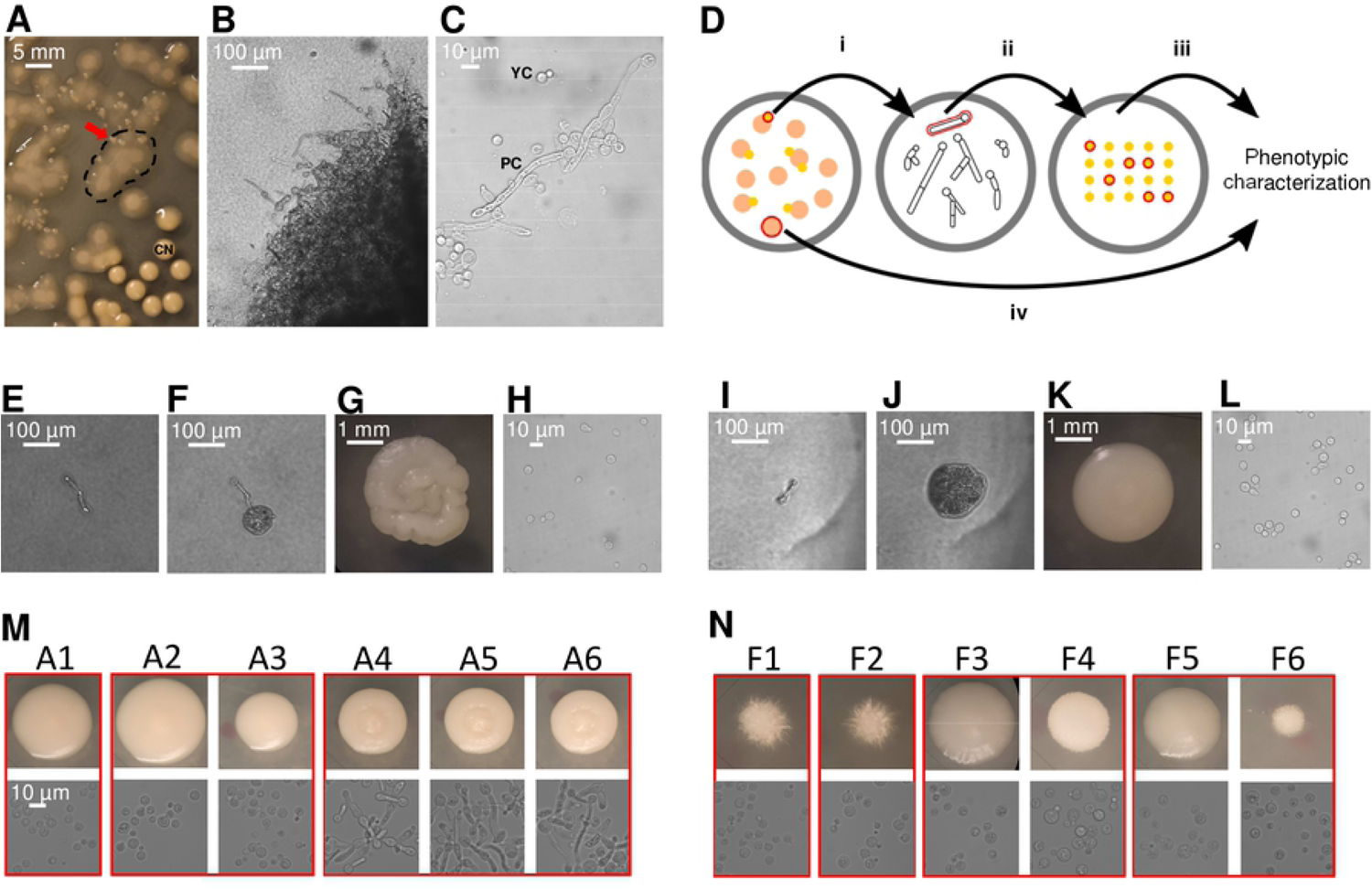
*C. neoformans* colonies exhibit various cellular and colony morphologies after co-incubated with amoebae in Sabouraud agar. (A) Small colonies of H99 (red arrow) surviving in a mat of amoeba that appears a hazy cloudy area (denoted by dashed line). Typical *C. neoformans* colonies (CN) are visible on the right bottom of the image. (B) Cells in the survival colony exhibit hyphal or pseudohyphal morphology (100× magnification). (C) Both pseudohyphae and yeast cells were identified on a wet mount of samples taken from the survival colony (400x). (D) Schematic representation of individual hyphae isolation. i. Survival colonies were picked using pipette tips and transferred to PBS in 3 mm culture dish. ii. Total 20 individual hyphae from 2-4 colonies were selected under microscope and transferred to fresh Sabouraud agar. Plates were incubated at 30 °C to generate colonies. iii. Six colonies were then selected for further phenotypic characterization. iv. Control colonies were also picked from the same plate of hyphal isolates but without interacting with amoeba. (E) Single pseudohyphal cell has been isolated from the survival colonies and transferred onto a fresh amoebae-free solid medium where form new colonies. (F) Microcolony with mostly yeast cells has been formed from a single pseudohyphal cell in 24 h. (G) Colony developed a serrated appearance after 2 days. (H) Yeast cells were identified on a wet mount of samples taken from the serrated colony. (I-L) Images showed another example of single pseudohyphal cell isolation. Smooth colony was formed from this particular pseudohyphal cell. (M-N) Same experiment was performed on environmental strains A1-35-8 and Ftc-555-1. Various cellular and colony morphologies have been identified among isolates A1-A6 and F1-F6 in the background of A1-35-8 and Ftc555-1 respectively. Colonies grew up from individual hyphae which was isolated from the same survival colony were grouped in red boxes.

Six colonies from each strain were selected together with three controls, which were colonies on the same plate with isolates but without interacting with amoeba, for further phenotypic characterization (Fig 1D). These will be referred heretofore to as amoeba-passaged isolates with numbers preceded by the letters H, A, and F to indicate their origin from strains H99, A1-35-8 and Ftc555-1, respectively. Controls will be referred to as C1-3 and ancestor will be referred to as A. To test if amoeba exerts selection pressure that resulted in amoeba-resistant cells, we examined if those isolates increased their survival during amoeba interaction. Isolates were then co-incubated with amoeba in the agar medium again, with *C. neoformans* in a cross, and amoeba were spotted in the center (Fig 2A). The radii of clear zones were measured as a function of time and these represented how well the amoeba clears the culture of *C. neoformans*. All of the amoeba-passaged isolates derived from H99 had reduced size of predator zone, when compared with their controls and ancestor strain (Fig 2B). In particular, the isolates that formed smooth colonies (H13, H16, H17) had the smallest predation zone (Fig 2B). This result implies that amoeba passage resulted in *C. neoformans* strains with increased ability to subsequently resist predation by amoeba. Next, we investigated the mechanism of the resistance. Samples were taken at the edge of the predator zone at the early stage of the interaction (week one), and observed under microscope. Isolates H13, H16 and H17 formed pseudohyphae while most of the cells in isolates H1, H2, H14 were in yeast form, with some displaying pseudohyphae (Fig 2C). However, no pseudohyphae were found in cells from controls and H99 ancestor colonies although pseudohyphae were formed eventually at the late stage of the interaction. Samples were also taken at a distance from the predation zone where cryptococcal cells had no contact with amoebae and in each of these regions all cells were in yeast form (Fig 2D). These results showed that pseudohyphal cells emerged rapidly from each of the amoeba-passaged strains even thought their cells were yeast prior to the incubation with amoebae and that pseudohyphal formation is a major mechanism of increased ability to resist predation.

**Fig 2.**
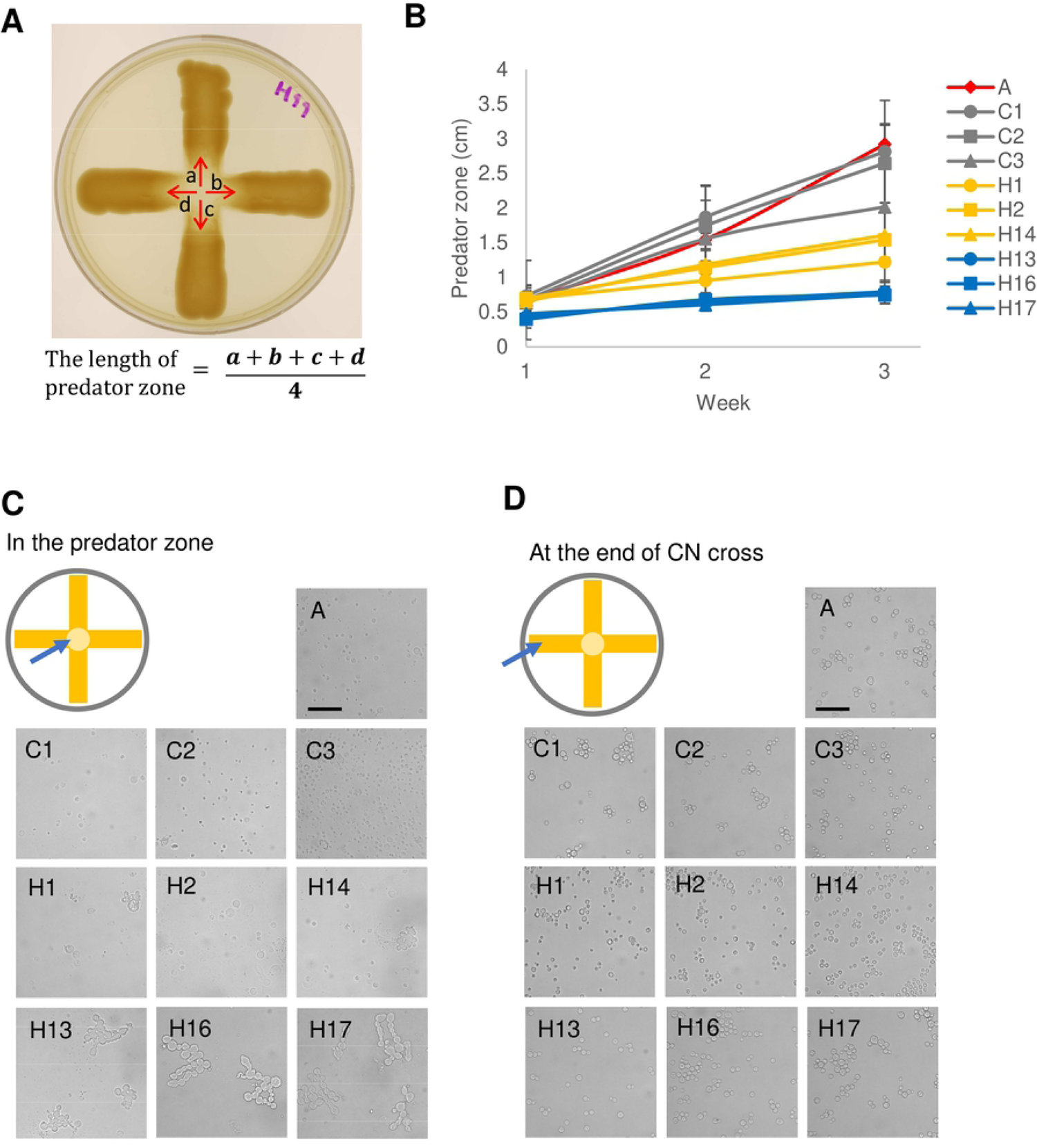
Isolates in H99 background derived from exposure to amoeba demonstrated increased resistance to amoebal killing by rapid pseudohyphal formation. (A) Scheme of amoebae killing assay. *C. neoformans* was streaked in a cross while *A. castellanii* was dropped at the intersection of the cross on Sabouraud agar. The data shown are the average of the distance between boundary and center of clear predator zone in four indicated directions (a-b), with the area being the predation zone. (B) All of the isolates that had prior exposure to amoeba had smaller clear zone than their ancestor and controls, consistent with enhanced resistance. A, ancestor; C1-3, controls; H, isolates derived from H99 after exposure to amoeba. Data are means from three biological replicates and error bars are SD. (C) Samples were taken from the peripheral areas of the predator zone after one-week co-incubation with amoebae and visualized under microscope. All of the isolates showed pseudohyphal formation, but ancestor and controls did not. (D) Sample were taken from the end of the cross where *C. neoformans* have not contacted with *A. castellanii* yet. All of the isolates manifested yeast cell morphology.

When the isolates derived from A1-35-8 and Ftc555-1 strains were again exposed to *A. castellanii*, some but not all exhibited increased resistance to amoebae (Fig 3A & B). Isolates derived from A1-35-8 (A4-6) were significantly more resistant than the others (Fig 3A). That may be due to maintenance of pseudohyphal cell morphology by isolates A4-6 even in the amoebae-free medium. Isolates F3-5 manifested increased resistance to amoeba but unlike the H99 derived isolates, displayed no pseudohyphal formation but had larger cells when compared to their ancestor (Fig 3C & D) at the early stage of interaction, which may be another survival strategy for *C. neoformans* against amoebae. In this regard, phagocytosis of *C. neoformans* by macrophage was reduced by cell enlargement of *C. neoformans* [49–51]. The resistance of isolates F3-F5 to amoebae may reflect their larger cell size.

**Fig 3.**
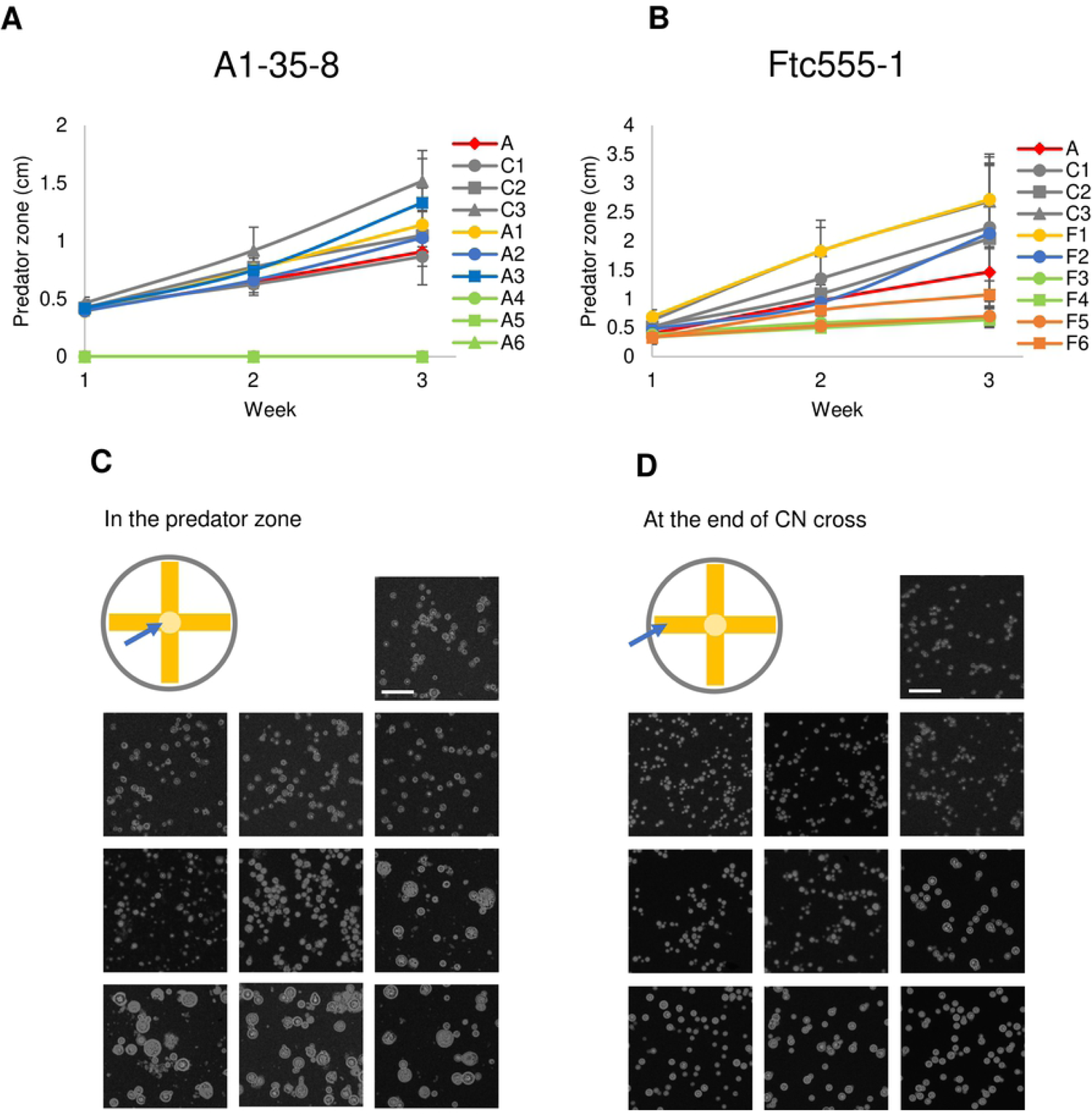
Some of the isolated recovered from the environmental strains A1-35-8 and Ftc-555-1 exhibited increased resistance to *A. castellanii*. (A) No clear predator zone of clearance was apparent with isolates A4-A6, while larger predator zones were apparent for isolates A1-A3 when comparing to their ancestor. A, ancestor; C1-3, controls; A1-6, isolates derived from A1-35-8 after exposure to amoeba (B) Isolates F3-F5 showed smaller predator zone than their ancestor. Data are means from three biological replicates and error bars are SD. A, ancestor; C1-3, controls; F1-6, isolates derived from Ftc555-1 after exposure to amoeba (C) Ftc-555-1 samples were collected from predator zone after one-week co-incubation. Isolates F3-F6 formed larger cell size than their ancestor and controls. (D) The cell size of isolates F3-F6 from the end of the cross is slightly larger than the one of ancestor and controls, but they are not as large as the cells taken from predator zone.

### Effects of amoeba selection on known virulence factors

*C. neoformans* expresses virulence factors that promote its pathogenicity, including formation and enlargement of a polysaccharide capsule, melanin production, extracellular secretion of urease, and cell enlargement. To evaluate whether the emergence of variant form of *C. neoformans* was accompanied by changes to known virulence factors, we analyzed the virulence-related phenotypic characteristics of the isolates derived from the three strains. Isolates H13, H16 and H17 had larger capsule thickness relative to their ancestor when cultured in minimal medium but cell sizes were similar (Fig 4A and 5A). All of the isolates also had increased urease activity in comparison to their ancestor (Fig 6). Isolates H1, H2 and H14 manifested less melanin production (Fig 7). We also examined if there were any changes of virulence factors in isolates of environmental strains. Each of the isolates derived from A1-35-8 strain had increased capsule size, reduced melanin production and increased urease activity when comparing to their ancestor, but there was no change in cell size (Fig 4B, 5B and 6). Isolates F3-F6 has increased both their capsule and cell size in minimal medium (Fig 4C and 5C). They were also having 15-18% of cells with size larger than 10 µm inside macrophages (Fig 5E) while approximately 80% of cells with larger than 10 µm in macrophage medium in 37°C 9.5% CO_2_ (Fig 5D). Moreover, all of the isolates of Ftc555-1 strain had increased urease activity, but reduced melanin production (Fig 6 and 7).

**Fig 4.**
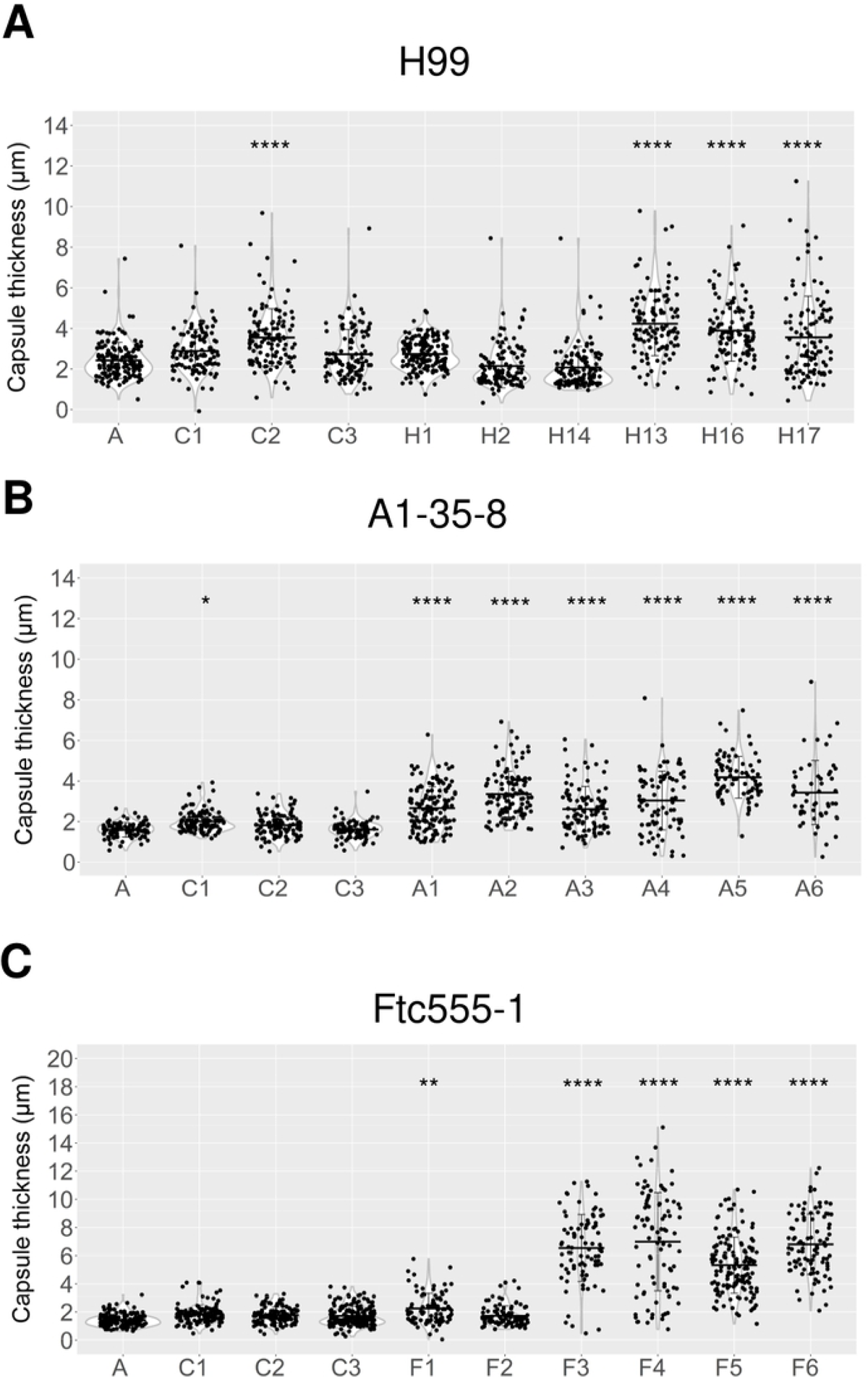
Capsule thickness for cells of parent strain and amoeba-selected strains. (A) H99 isolates (B) A1-35-8 isolates and (C) Ftc555-1 isolates have been cultured in minimal medium at 30 °C for three days. Capsule was visualized by counterstaining with India ink. A, ancestor; C1-3, controls. * *P* < 0.1 ** *P* < 0.01 **** *P* < 0.0001 by One-way ANOVA, followed by Tukey’s multiple-comparison test.

**Fig 5.**
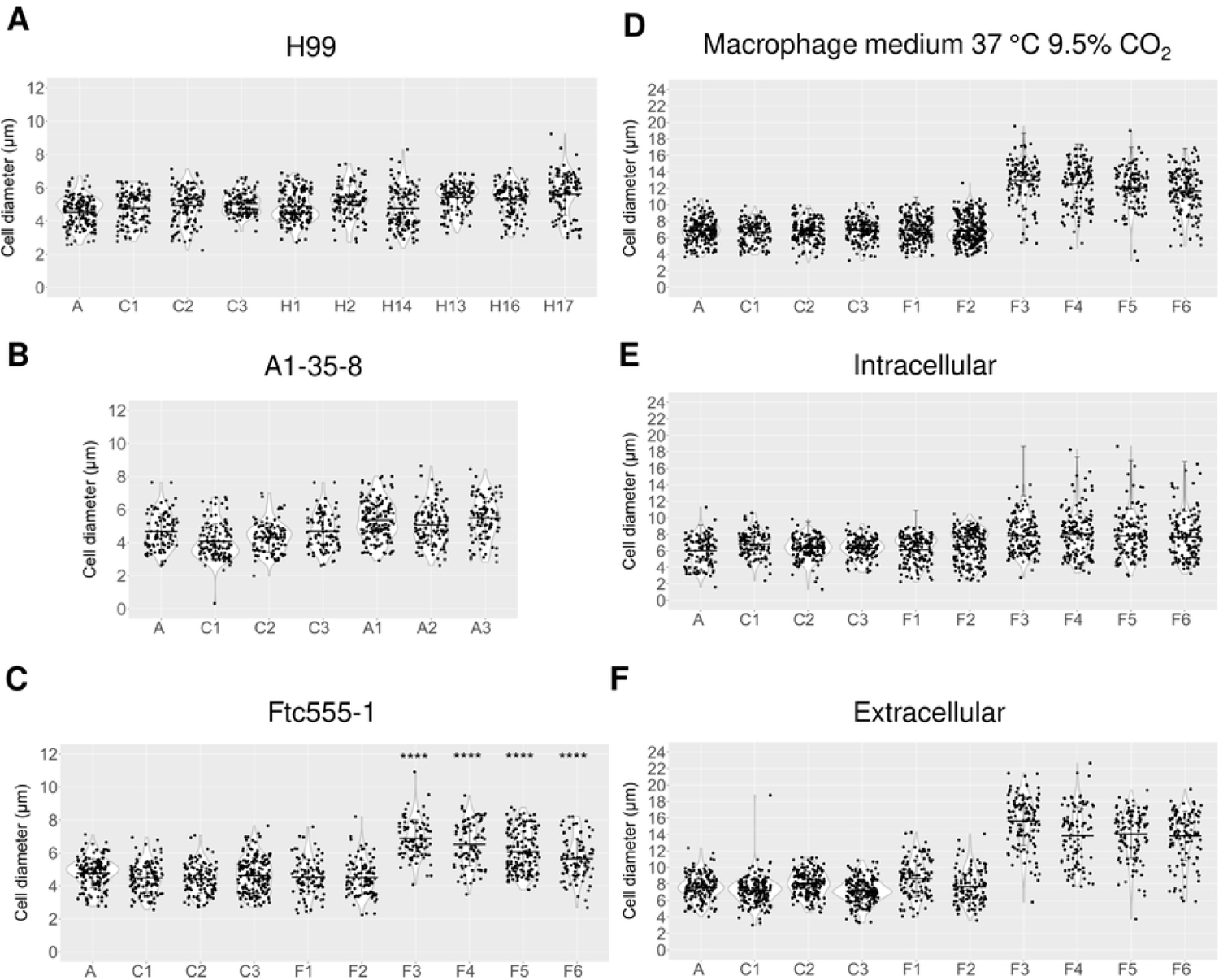
Cellular dimensions for cells of parent strain and amoeba-selected strains. (A) H99 isolates (B) A1-35-8 isolates and (C) Ftc555-1 isolates have been cultured in minimal medium for three days. A, ancestor; C1-3, controls. (D-E) Ftc555-1 isolates have also been cultured in macrophage medium and with BMDM at 37 °C 9.5% CO_2_ for 24 h. Extracellular cryptococcal cells were collected from the culture supernatant while intracellular cells were retrieved from lysing the BMDM. A, ancestor; C1-3, controls. **** *P* < 0.0001 by One-way ANOVA, followed by Tukey’s multiple-comparison test.

**Fig 6.**
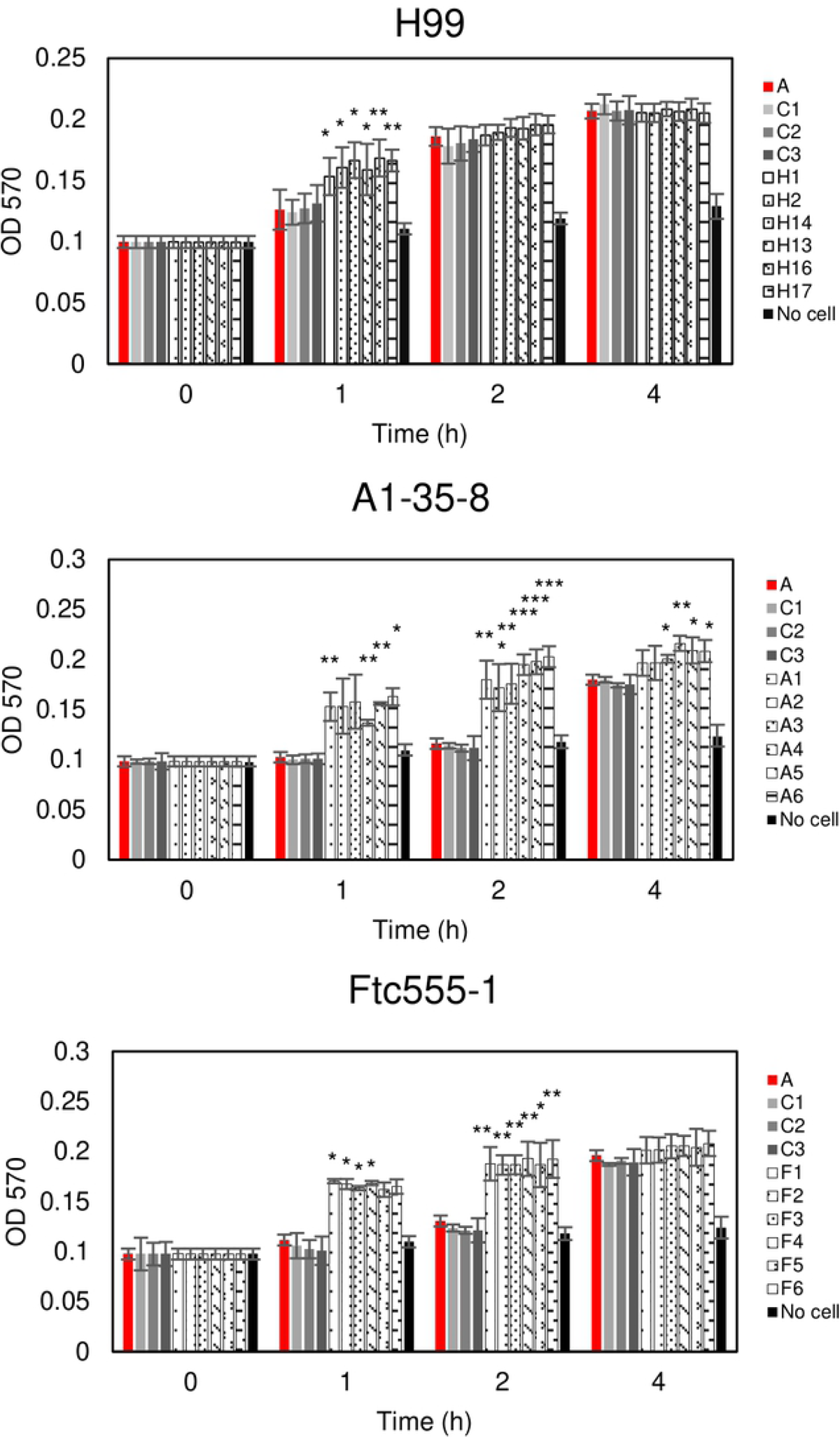
Urease activity for cells of parent strain and amoeba-selected strains. The urease activity of cryptococcal cells were detected by using rapid urea broth (RUH) method. Amoeba-passaged isolates with numbers preceded by the letters H, A, and F to indicate their origin from strains H99, A1-35-8 and Ftc555-1, respectively. A, ancestor; C1-3, controls. The assay was performed in triplicate for each time point. Error bars represent SD. * *P* < 0.1 ** *P* < 0.01 *** *P* < 0.001 by unpaired t-test.

**Fig 7.**
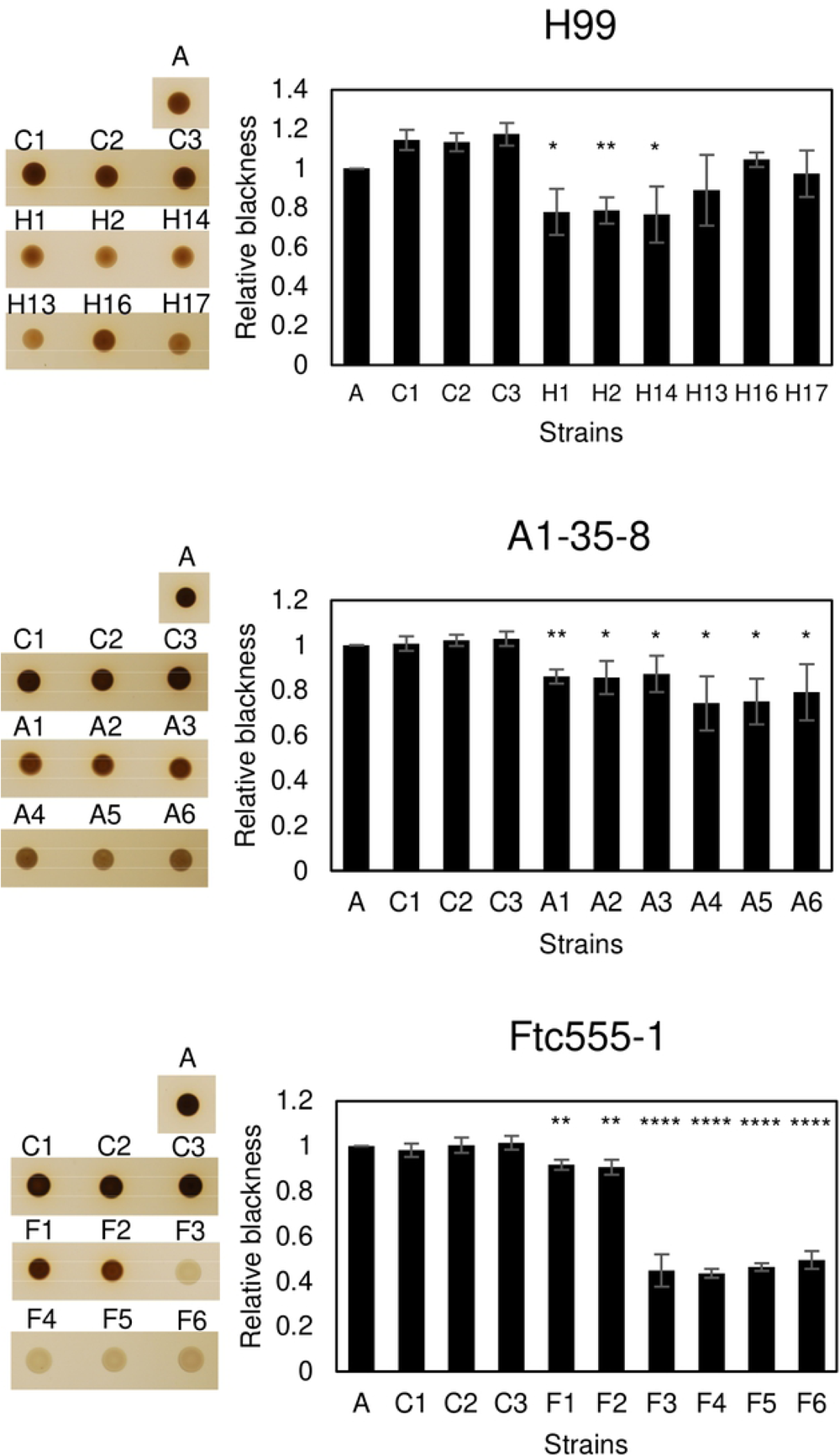
Melanization formation for cells of parent strain and amoeba-selected strains. Melanization was analyzed by spotting the 10^6^ cryptococcal cells on minimal medium agar with L-DOPA for 24 h. The pigmentation of colony was measured through grayscale pixel quantification by the software ImageJ. Relative blackness was calculated as a ratio of grayscale quantification between isolates, their ancestor (A) and control (C1-3). Error bars represent SD. * *P* < 0.1 ** *P* < 0.01 **** *P* < 0.0001 by unpaired t-test.

We further characterized the isolates in stress conditions by analyzing their growth under thermal stress and exposure to the antifungal drug fluconazole (Fig 8). Isolates H13, H16, H17 had reduced growth at 40°C and in the presence of fluconazole while H1, H2 and H14 had slightly increased their resistance to fluconazole compared to their ancestral strain. Isolates A4-A6 and F3-F6 displayed defects in growth at high temperature and after exposure to fluconazole. Overall, the data show that the phenotypic changes were broad and diverse among isolates.

**Fig 8.**
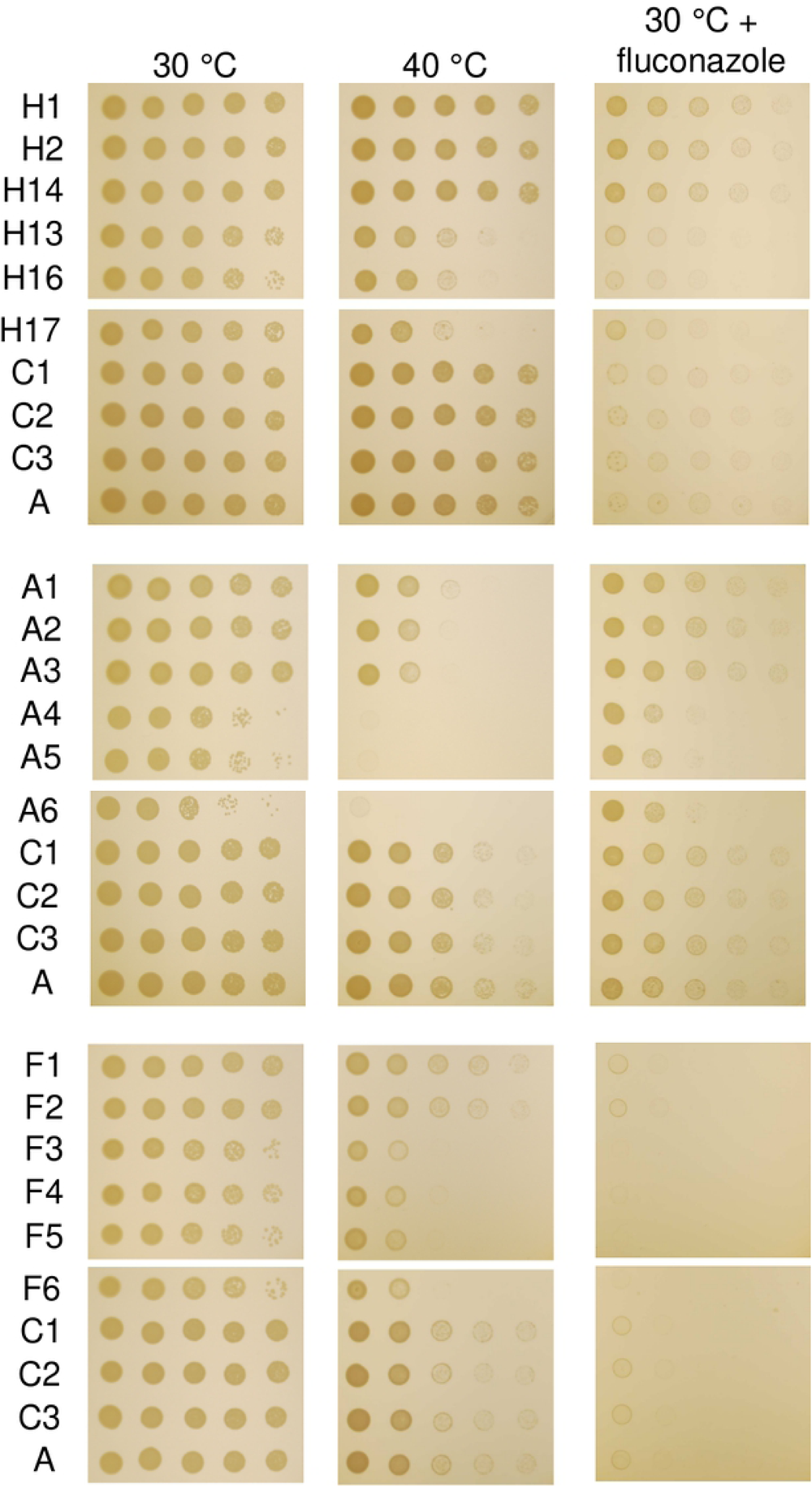
The growth of parents and isolates under stress conditions. Cells were 10-fold serially diluted and spotted onto YPD medium with or without fluconazole (16 µg/ml), and grown for two days at 30°C or 40°C. Amoeba-passaged isolates with numbers preceded by the letters H, A, and F to indicate their origin from strains H99, A1-35-8 and Ftc555-1, respectively. A, ancestor; C1-3, controls.

### Genomic analysis and sequencing results

A prior study showed that DNA mutation was involved in pseudohyphal formation during amoebae interaction [52]. To find out if there are any such mutations or any other mutations in our experiments, the genomes of all isolates were sequenced. SNPs and indels were identified compared to the H99 reference genome (Table 1, 2 and table in S1 table). Genome sequencing revealed that H and A isolates acquired only a small number of SNPs and indels during amoeba passage, whereas F isolates acquired an order of magnitude more SNPs and indels. Two SNPs were identified in H1, H2, H14 in comparison with their ancestral strain H99. One of the SNPs is a missense mutation (M484R) in a gene encoding an oligopeptide transporter (CNAG_03013; *OPT1*). This mutation creates the replacement of methionine 484 with arginine. Opt1 has been shown to be required for transporting Qsp1, a quorum sensing peptide, into the receiving cells [53]. Another SNP is an intron variant in a gene encoding a protein kinase (CNAG_02531; *CPK2*) as part of the MAPK protein kinase family. Loss of CPK2 reduces melanin production in Niger seed media. For A1-35-8 derived isolates, a total of four SNPs were identified relative to the ancestral strain A1-35-8. In A1, one missense SNP was found in CNAG_01101, which encodes a hypothetical protein with a centrosomin N-terminal motif and also a single nucleotide deletion was identified in CNAG_03013, causing a frameshift at P358. Two SNPs were identified in A2 and A3 isolates, with one SNP leading to nonsense mutation in CNAG_03013 and another SNP resulting in missense mutation in CNAG_02858 which encodes adenylsuccinate synthetase. Another SNP in the A2 isolate was found in an intergenic region, a site with a high fraction of ambiguous calls. Isolates A4-6 had a single nucleotide deletion at gene CNAG_03622 (*TAO3*) and leads to the frameshift at residue 150 of 2392. This mutation is consistent to the finding in previous study that *TAO3* mutation lead to the pseudohyphal phenotype [52]. In contrast to the A1-35-8 isolates, the rate of mutations in the Ftc555-1 isolates was 10 times higher, ranging from 22 to 77 SNPs (total 225 SNPs) and 7-15 indels (total 34 indels) in these isolates. Among those SNPs, three SNPs were annotated as high impact mutations resulting in disruption of the coding region (early stop codons and splice site mutations). One SNP results in a nonsense mutation (G407*) in CNAG_00570 which encodes Pkr1 (AMP-dependent protein kinase regulator) in F5 and F6 isolates. In addition, F3 and F4 isolates carry a single nucleotide deletion in CNAG_00570 leading to frameshift of residues 194 of 482. Pkr1 is one of the important components of cAMP/PKA pathway and negatively regulates Pka activity which is involved in morphogenesis, nutrient acquisition, stress responses and virulence in *C. neoformans* (Choi et al., 2012). Another SNP in the F1 isolate is a splice site mutation in CNAG_03013. In summary, there are three noteworthy observations in the sequence data: 1. The gene CNAG_03013 (*OPT1*) was impacted by non-synonymous SNP changes in all three strain backgrounds; 2. The previously described *TAO3* mutation responsible for pseudohyphal or hyphal formation was found in our isolates A4, A5, A6 [52]; and, 3. No SNPs and indels were found in some of the isolates including H13, H16, H17 suggesting that the phenotypic changes observed did not originate from single nucleotide variants in the genome.

**Table 1.** High and moderate impact SNPs found in passaged isolates

| Isolates | Chr | Position | Reference | Alternate | Gene ID | Gene function | Effect of mutation |
| --- | --- | --- | --- | --- | --- | --- | --- |
| H1, H2, H14 | 3 | 211613 | T | G | CNAG_03013 | Oligopeptide transporter | M484R |
| H1, H2, H14 | 6 | 68953 | C | A | CNAG_02531 | Calcium-dependent protein kinase | Intron variant |
| A1 | 5 | 1208219 | T | C | CNAG_01101 | Hypothetical protein | R478G |
| A2, A3 | 3 | 213165 | G | A | CNAG_03013 | Oligopeptide transporter | Nonsense mutation W932* |
| A2, A3 | 3 | 594765 | A | G | CNAG_02858 | Adenylsuccinate synthetase | I346V |
| A2 | 13 | 592173 | C | T |  | Intergenic region |  |
| F1<sup>†</sup> | 3 | 213566 | G | T | CNAG_03013 | Oligopeptide transporter | Splice site mutation |
| F5, F6<sup>†</sup> | 1 | 1469244 | C | A | CNAG_00570 | cAMP-dependent protein kinase regulator | Nonsense mutation G407* |
| FC2<sup>†</sup> | 11 | 136455 | T | G | CNAG_01506 | Hypothetical protein | Splice site mutation |

<sup>†</sup> Only high impact mutations of Ftc555-1 variants were shown in this table.

**Table 2.** High impact indels found in passaged A1-35-8 isolates

| Isolates | Chr | Position | Reference | Alternate | Gene ID | Gene function | Effect of mutation |
| --- | --- | --- | --- | --- | --- | --- | --- |
| A1 | 3 | 211137 | GC | G | CNAG_03013 | Oligopeptide transporter | Frameshift at P358 |
| A4, A5, A6 | 2 | 363200 | CA | C | CNAG_03622 | Cell polarity | Frameshift at N150 |

Table 2 High impact indels found in passaged A1-35-8 isolates
| Isolates | Chr | Position | Reference | Alternate | Gene ID | Gene function | Effect of mutation |
| --- | --- | --- | --- | --- | --- | --- | --- |
| A1 | 3 | 211137 | GC | G | CNAG_03013 | Oligopeptide transporter | Frameshift at P358 |
| A4, A5, A6 | 2 | 363200 | CA | C | CNAG_03622 | Cell polarity | Frameshift at N150 |

To determine if the high impact mutations we identified in genes *PKR1, OPT1*, CNAG_02531 and CNAG_01506 are responsible for resistance to the killing of amoebae, deletion mutants of the candidate genes in the H99 background were co-incubated with amoebae on solid medium. However, the predator zones from these mutants were comparable with the parental strain (S2 Fig).

### Aneuploidy

We next hypothesized that emergence of aneuploidy could be a source of evolutionary adaptation because aneuploidies are frequent in *C. neoformans* and it has been shown to play crucial roles in stress resistance [54,55]. To this end, the chromosomal copy numbers of the isolates were defined based on the normalized depth of sequence coverage. The analysis revealed that there were duplications of chromosome 8 in isolates H13, H16 and H17, but no chromosomal duplication has been found in other isolates (Fig 9A). The results were confirmed by qPCR with two selected isolates, H14 and H17 (Fig 9B and S1 Fig). We next investigated if this chromosomal duplication was responsible for the pseudohyphal formation and other phenotypic changes. In order to do so, H17 was passaged in fresh rich medium every day for 30 days to eliminate the duplication. The elimination was confirmed by qPCR (Fig 9C). H17 euploid strain (H17^eu^) was then co-incubated with amoebae culture in solid medium, and samples were taken from the edge of the predation zone and visualized under microscope. No pseudohyphae could be observed in H17^eu^ (Fig 9D). In such case, the observation was similar to what we found in H99, but distinct from H17 aneuploid strain (H17^aneu^) that forms mostly pseudohyphae after one-week co-incubation (Fig 9D). Not surprisingly, H17^eu^ had decreased ability for amoebal resistance, having a similar size of the predator zone as H99 while H17^aneu^ had a smaller predator zone (Fig 9E). The capsule size of H17^eu^ was smaller than H17^aneu^ and similar to H99, suggesting that the duplication of chromosome 8 results in larger capsule size (Fig 9F). H17^eu^ had lower urease activity than H17^aneu^ but comparable level with H99 after 1 h (Fig 9G). However, the urease activity of H17^eu^ increased faster than that of H99 after 1.5 h. The result implied that the chromosomal duplication may be responsible partially the high urease activity found in H17^aneu^.

**Fig 9.**
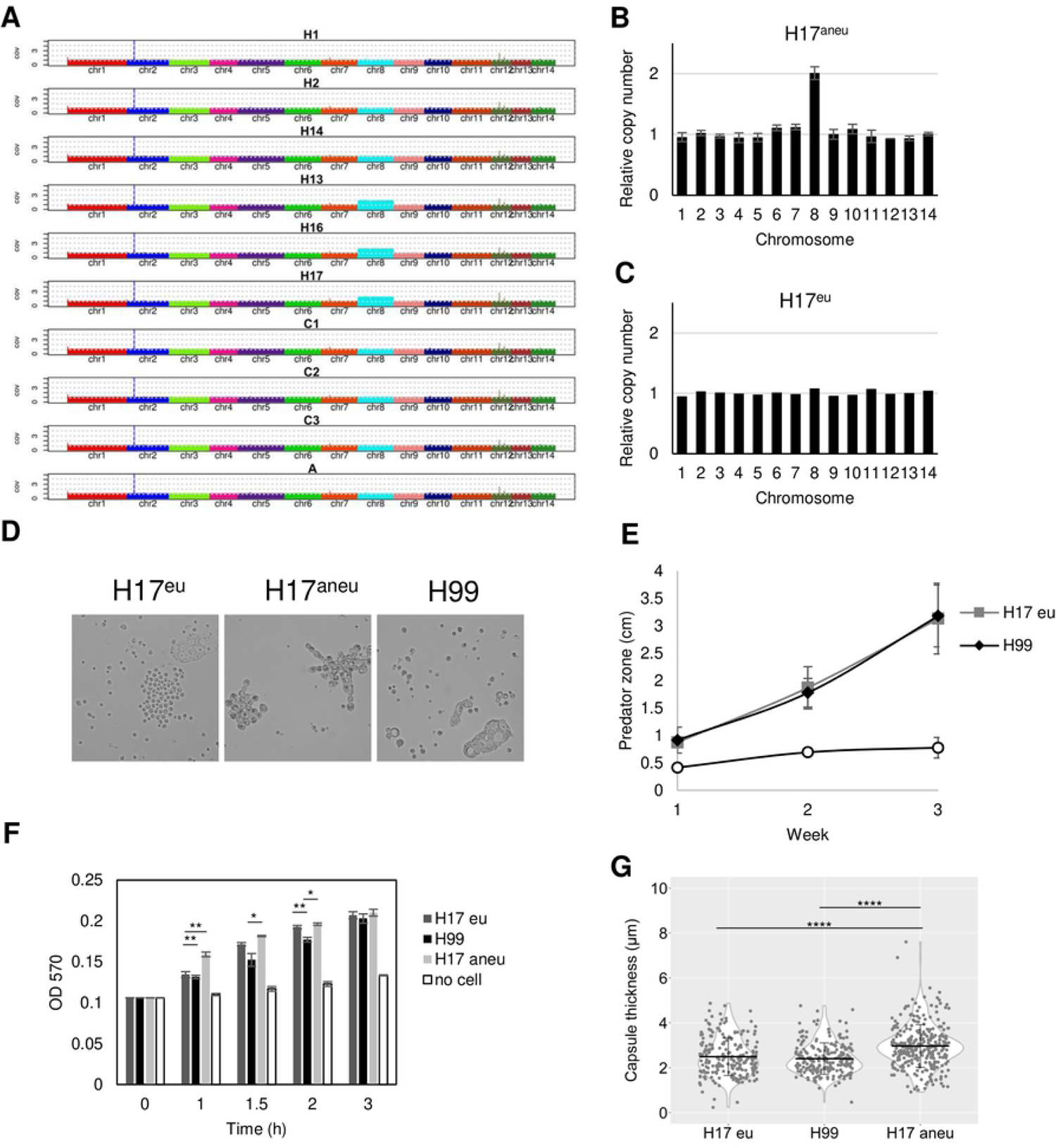
Aneuploidy plays a role in pseudohyphal formation. (A) Chromosomal copy numbers of H99 isolates were determined based on depth of sequence coverage normalized by the average genome-wide sequence depth (B) Relative chromosome copy number of isolate H17 was obtained by qPCR. H17 have duplication of chromosome 8 (C) chromosome duplication in H17 is eliminated by passaging H17 in fresh Sabouraud medium for 30 days. (D) H17 euploid (H17^Eu^) strain did not form pseudohyphae as rapid as H17 aneuploid strain. (E) H17^Eu^ euploid strain has larger predator zone than H17^Aneu^. Data represent the mean of three biological replicates per biological sample and error bars are SD. (F) H17^Eu^ strain has lower urease activity then H17 ^Aneu^ and comparable urease activity as H99 at early time point (1 h) Data represent the mean of two biological replicates per biological sample and error bars are SD. (G) H17^Eu^ has smaller capsule size than H17^Aneu^, but similar capsule size with H99.

Aneuploidy can arise from multinucleate state through transient polyploidization after failed cytokinesis or cell fusion. The filamentous multinucleate fungus *Ashbya gosypii* exhibits both polyploidy and aneuploidy frequently after cell division [56]. Since pseudohyphae have a cytokinesis defect and multinuclei within a common cytosol, we asked if the pseudohyphal formation may lead to ploidy variation and thus may become one of the sources of phenotypic variation. Consequently, H99 expressing green fluorescent protein-labeled histione-2 (GFP-H2B) were visualized by time-lapsed imaging, and a nucleus fusion have been observed in one of the pseudohyphae after nuclei separation (Fig 10 & S1 Movie). This event provides evidence that polyploidization can exist in pseudohyphae and thus may have a high chance of leading to aneuploidy and phenotypic variation.

**Fig 10.**
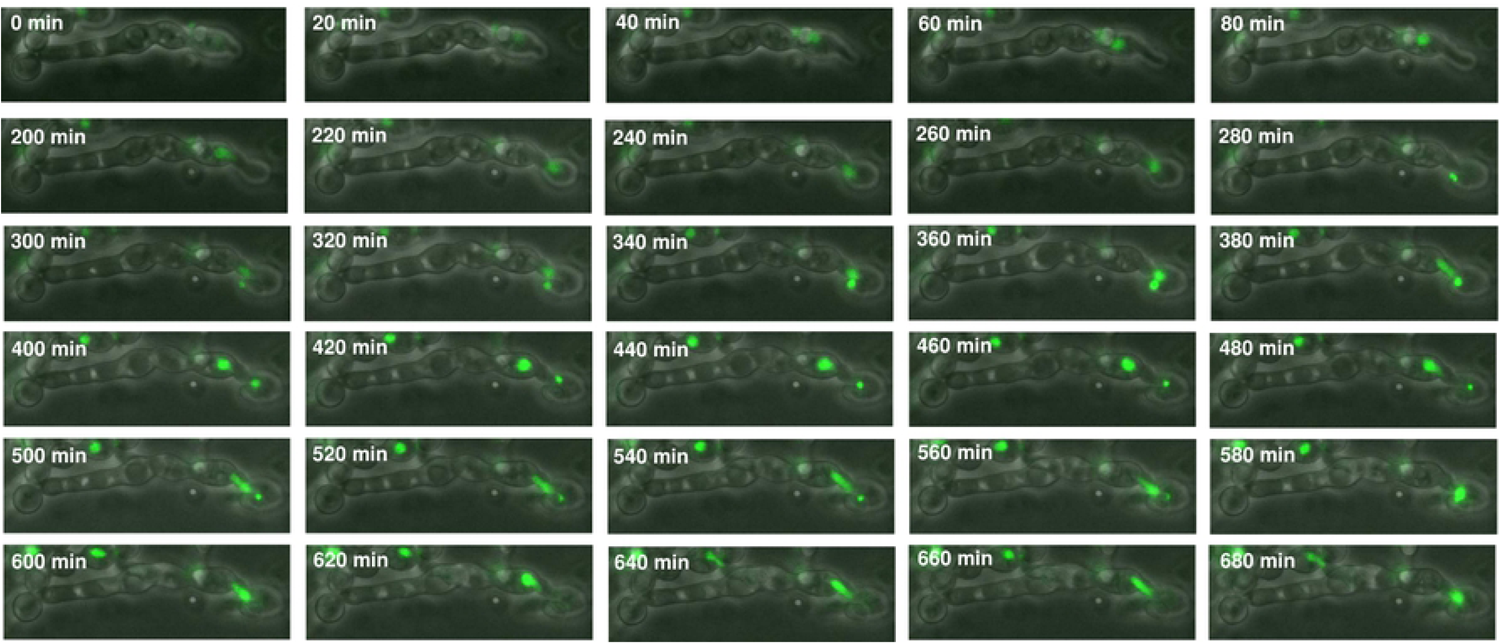
Time-lapse imaging showing nuclear division of pseudohyphae. The images of pseudohyphae of amoeba-passaged H99 GFP-H2B strain were taken by phase-contrast and fluorescence microscopy. Bud (red arrow) were forming between 0-220 min. The nucleus migrates into the daughter cells at 240 min, and separated at 300 min. Nuclear division was completed at 400 min. However, the nucleus from mother cells re-entered into the daughter cells at 500 min and underwent fusion at 580 min.

### Epigenetic modifications

Chromatin remodeling can rapidly moderate transcriptional response in order to let the microorganism to adapt rapidly to stressful conditions in hosts. Histone acetyltransferase activity has been shown to be essential for *C. neoformans* in virulence regulation and response to host environments [57]. Since the phenotypes changed rapidly after *C. neoformans* interacted with amoebae, we hypothesized that histone acetylation may be involved in the phenotypic changes. To determine whether the evolution of *C. neoformans* after interacting with amoebae also involved histone modification, we compare the quantitation of the acetylation of core histone H3. However, we did not detect significant differences in global histone H3 acetylation between isolates and ancestral strains (S3 Fig).

### Effects of amoeba selection on interactions with murine macrophages

Based on the changes of multiple virulence-related phenotypes, we expected that some of the isolates would have a better survival when interacting with macrophages. However, there was no significant change of intracellular survival among all the isolates (Fig 11A-C). Nevertheless, we cannot rule out the possibility that isolates may cause damage to macrophages. Since isolates F3-F6 underwent cell enlargement inside macrophage, we hypothesis the increased cell size may physically rupture macrophages. Therefore, we measured the release of lactate dehydrogenase (LDH) from the macrophage when they were infected with Ftc555-1 isolates. Indeed, it was found that LDH release was significantly induced from the macrophages containing F3-F6 when compared to the ancestral strain (Fig 11D), suggesting that F3-F6 and their enlarged yeast cells cause certain damages to their host cells.

**Fig 11.**
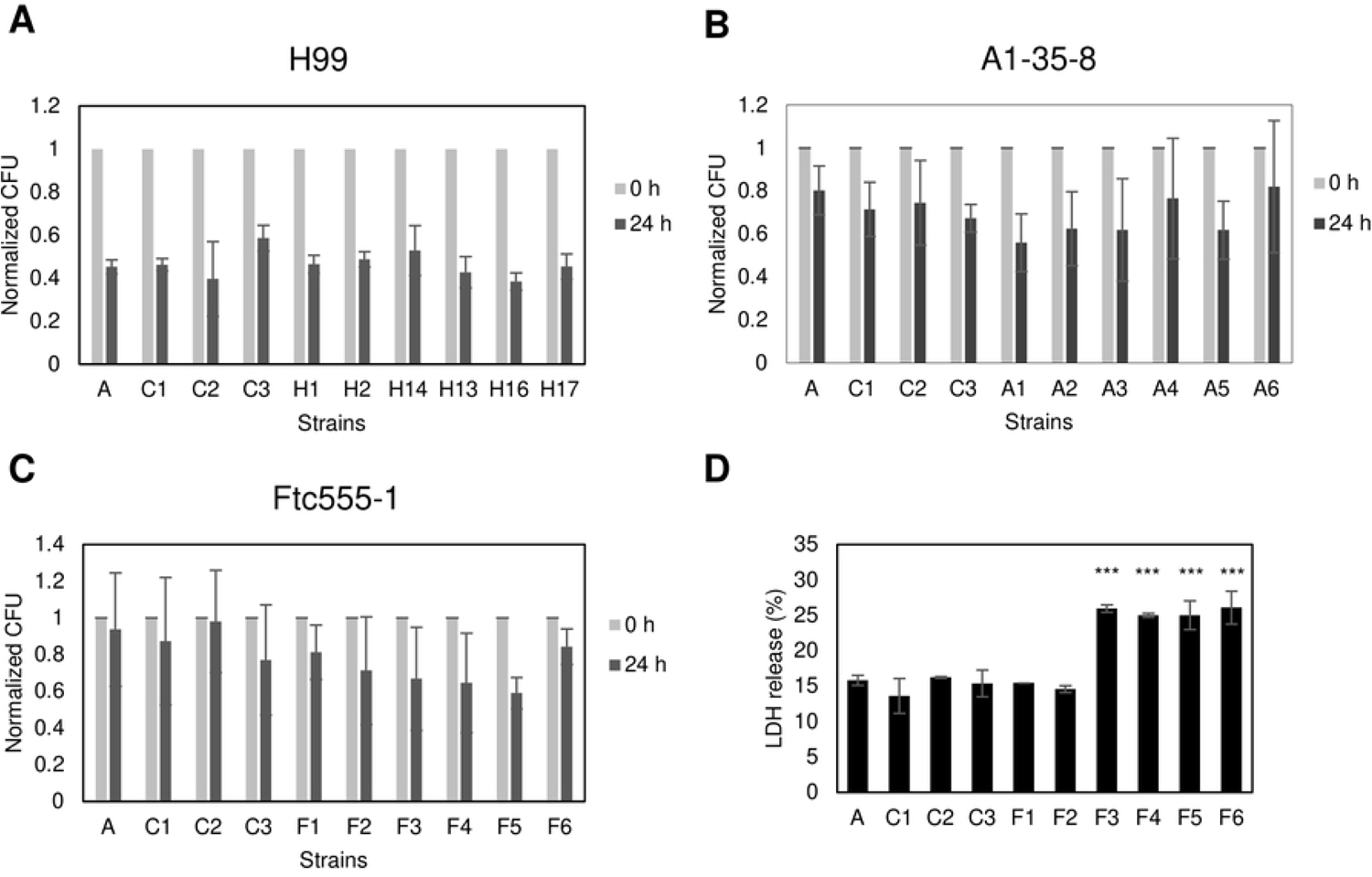
(A-C) The survival of parents and isolates in culture with BMDM. The survival of (A) H99, (B) A1-35-8 and (C) Ftc555-1 isolates was determined by colony form unit (CFU) after 0 and 24 h phagocytosis. The percentage of survival was calculated by normalizing the CFU value of 24 h infection to time zero. A, ancestor; C1-3, controls. Data represent the mean of three biological replicates and error bars are SD. (D) BMDM were infected with Ftc555-1 isolates for 48 h. LDH release from damaged BMDM into culture supernatant was assayed. *** *P* < 0.001 by One-way ANOVA, followed by Tukey’s multiple-comparison test.

### Virulence testing in murine model and moth larvae. The deletion of PKR1 has been reported to be manifest hypervirulence in mice infection [58]

Since isolates F5 and F6 were more cytotoxic to macrophages and contained loss of function mutations in *PKR1*, we investigated the virulence of F5 and F6 and their parental strain Ftc555-1 in a murine infection model. However, all animals survived after intranasal inoculation for 60 days (data not shown). Lung fungal burden was determined by enumerating CFU. Only the cells of the initial isolate (Ftc555-1) were detected in the mouse lung after this incubation period and there was considerable mouse-to-mouse variation in CFU. Hence, the two isolates carrying *PKR1* mutations were cleared from the lungs 60 days after inoculation (Fig 12). Consequently, we explored early times of infection and noted that at day 5 after challenge both Ftc555-1 and F5 had comparable fungal burden while that of F6 was reduced (Fig 12). To determine if the different proliferation rates of the strains are the reason of the fungal burden difference, proliferation analysis were performed to observe the growth curves of the three *C. neoformans* strains. Two growth curves were prepared in similar conditions, except the initial number of yeasts. Starting the curve with a high concentration of cells (1.0×10^7^) displayed a similar growth increase for F5 and F6 cells during the first 36 hours, but the very opposite was observed when the curve started with a low concentration of cells (5.0×10^3^), with a higher growth increase for Ftc555-1 strain (S4 Fig). Hence, all three strains were able to establish themselves in mice initially and survive clearance by innate immunity but the F5 and F6 were subsequently cleared, presumably by the development of acquired immunity.

**Fig 12.**
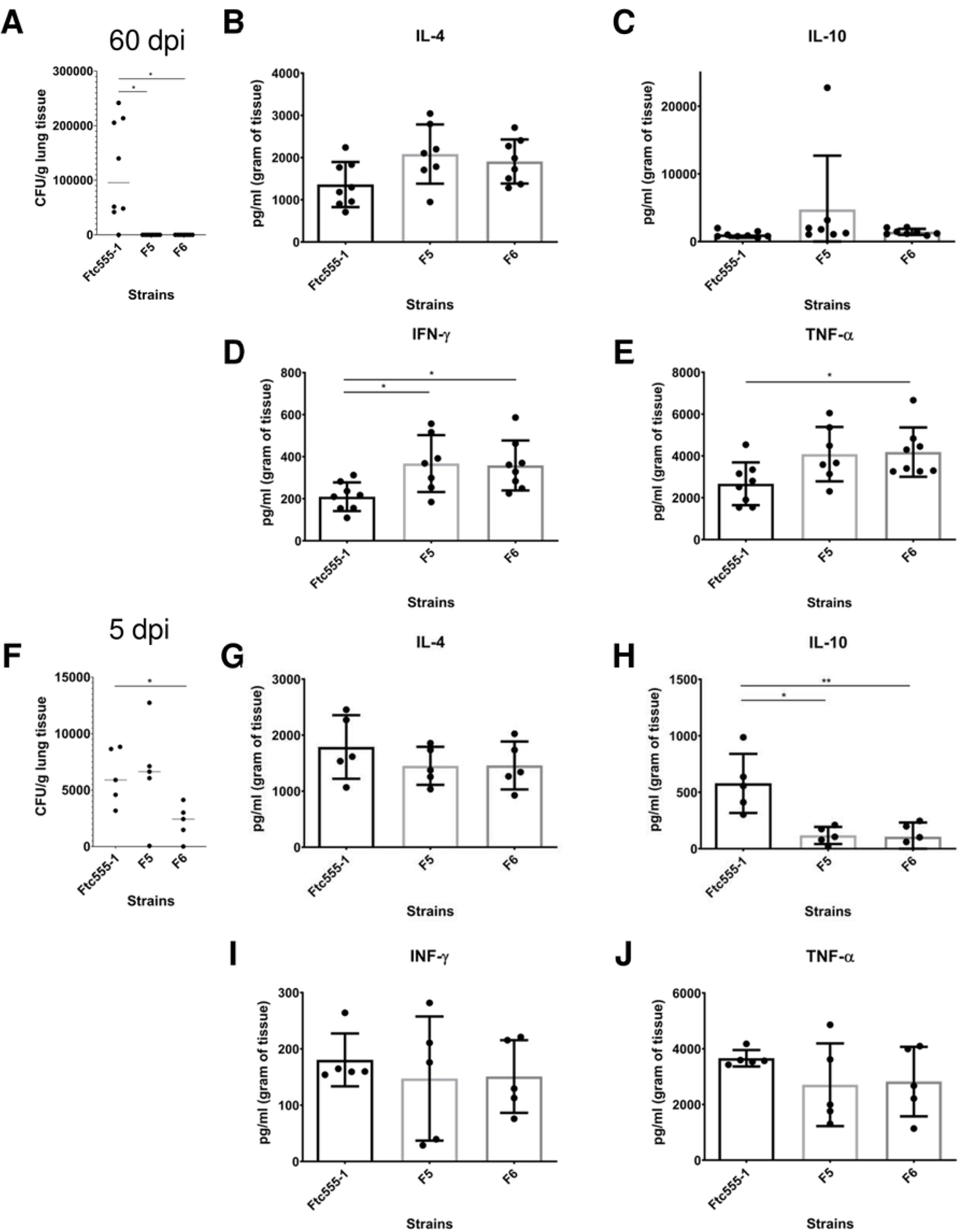
Fungal burden and cytokine production in lung after infection with Ftc555-1 as well as its isolates F5 and F6. After 60 days of infection, mice were sacrificed and (A) fungal burden in the lung were determined by CFU counting. The level of cytokines (B) IL4 (C) IL10 (D) IFN-γ (E) TNF-α in the lung were measured by ELISA. At 5-day post-infection, (F) fungal burden, and the amount of cytokines (G) IL4 (H) IL10 (I) IFN-γ (J) TNF-α in the lung were also measured. All data represent the mean of eight mice per group and errors bars are SD. * *P* < 0.1 ** *P* < 0.01. For determination of cytokine levels, one-way ANOVA with Kruskal-Wallis nonparametric test was used and followed by Bonferroni’s multiple-comparison test. The t-test was used to compare the number of colony forming units (CFU) for different groups.

To gain more insight into the immune responses elicited by Ftc555-1, F5 and F6 in the lung we studied several cytokine responses. In the lungs, the levels of pro-inflammatory cytokines, including tumor necrosis factor alpha (TNF-α) and interferon gamma (IFN-γ), were increased and remained high after the 60 d post-infection with F5 and F6 strains (Fig 12). The lung cytokine response shows that F5 and F6 elicit lower levels of IL-10 than Ftc555-1, which could help their clearance from lung tissue relative to Ftc555-1 since reduction of this anti-inflammatory cytokine has been associated with increased resistance to cryptococcal infection in mice [59]. Interestingly, the levels of the same molecules were different when we analyzed the systemic immune response as measured by cytokines in their spleens (S5 Fig). These results may indicate that sustaining high levels of these cytokines may stimulate an inflammatory reaction, which could be associated with resolution of the infection of the mutant strains. These cytokine results show that Ftc555-1 is eliciting quantitatively different immune responses from the parent strain Ftc555-1 consistent with the notion that the differences in virulence observed for these strains reflect differences in the effectiveness of the immune responses triggered.

We also examine the virulence of the isolates using wax moth larvae model, and isolates H13, H16 and H17 were less virulence than their ancestor (S6A Fig). This may due to the fact that these isolates can form pseudohyphae rapidly in the larvae, and pseudohyphal *C. neoformans* are attenuated for virulence in wax moth larvae [52]. However, so far, there was no statistically significant increased virulence of isolates in the wax moth larvae infection model (S6 Fig).

## Discussion

In the past two decades, the concept that amoeba acts as selective pressure for virulence traits of environmental microbes has gained considerable traction. For fungal pathogens, concordance between virulence factor function in amoeba and macrophages has been demonstrated for *C. neoformans* [14,40], *Aspergillus fumigatus* [23,60] and *Paracoccidioides spp*. [24], but many questions remain on how fungal-protozoal interactions select for mammalian virulence. In this study, we investigated how interactions with amoebae affected the phenotype and genotype of *C. neoformans* to explore the mechanisms behind this long-term evolutionary adaptation. Our results provide new insights on how amoeba predation can drive the evolution of *C. neoformans* since survivors emerge that show major phenotypic and genetic differences from the founder strain. The phenotypic diversity may facilitate *C. neoformans* adaptation to different hosts and thus enhance its virulence.

Pseudohyphae formation was the most common response to *C. neoformans* survival when faced with amoeba predation. This result confirms an older observation that pseudohyphal formation was a ‘escape hatch’ for *C. neoformans* survival when preyed upon by amoebae [61]. Different fungal morphologies are reported to trigger different killing mechanisms by amoeba [62] and the *C. neoformans* filamentous form may be more resistant to killing. Similar to our observation, Nielson et al [61] reported that when *C. neoformans* was co-cultured with amoebae, most of the fungal cells were killed with survivors forming colonies that contained pseudohyphae. Most of their isolates remained pseudohyphal, with only one out of eight isolates reverting back to the yeast form. That result differed from ours, since most of the pseudohyphal isolates in this study reverted to yeast forms after removal from the amoebae culture, such that only 3 of 18 isolates studied in detail maintained a stable pseudohyphal phenotype. Those three isolates (A4 – A6) have a single nucleotide deletion in *TAO3* gene shown in whole genomic sequencing, and consistent to mutations in RAM/MOR pathway ofthe pseudohyphal variants reported in the previous study [52].

Previous studies have focused primarily on cryptococcal isolates with pseudohyphae phenotypes derived from amoeba, but in this study, we investigated in detail those amoeba-resistant isolates with unstable pseudohyphal phenotypes. We found that although some of the isolates (H13. H16, H17) reverted to yeast, they were able to form pseudohyphae quicker than their parental strain when they were exposed to amoebae again. These isolates were less virulent in *Galleria* infection model, a finding consistent with prior reports that the pseudohyphae strains were less virulent in animal models. Interaction with amoebae also resulted in measurable virulence-related phenotypic changes in *C. neoformans*, confirming that amoebae can play a powerful role in the selection of virulence factors, which are related to the pathogenesis of human disease. Of note, we selected only six isolates from each strain for further characterization, but all of them had changes, suggesting that the microevolution occurs frequently and rapidly when exposed to amoebae. Moreover, the changes were pleiotropic and included differences in colony morphology, capsule size, cell size, urease activity, melanin production, susceptibility to thermal stress and an antifungal drug. However, isolates studied revealed a different configuration of phenotypic changes although they tended to cluster in groups from the same survival pseudohyphal colony (S7 Fig). Overall, the interaction of *C. neoformans* with amoebae-passaged isolates with increased phenotype diversity. Since there are many types of amoeboid predators in the soil and *C. neoformans* does not know the identity of the phagocytic predator, generating great diversity in strains could provide this fungus with insurance that some will survive. Hence, the diversity observed among isolates that survived amoeba predation suggests a bet hedging strategy for survival based on the generation of phenotypic diversity.

To identify the mechanism for the phenotypic changes, we compared the whole genome sequencing of isolates and ancestral strains using deep sequencing to identify point mutations, amplification or deletion of chromosomal segments and whole-chromosome aneuploidy. We found that there were only two SNPs in H99 derived isolates, four SNPs and two indels in the A1-35-8 derived isolates. Isolates from the same survival pseudohyphal colonies had similar SNPs, which is consistent with the similarity of their phenotypic changes, suggesting that the point mutations may be associated with some of the phenotypic changes. Interestingly, there were total 252 SNPs in Ftc555-1 derived isolates with an average of 48 SNPs among isolates (range of 22-80), a rate approximately10 times higher than H99 and A1-35-8. That may be explained by the fact that the ancestral Ftc555-1 strain contains a splice donor site mutation in *MLH1*, a gene involved in mismatch repair of nuclear DNA. This predicted high impact, loss of function mutation (G to A change at position 1270268 of chr 6) is also found in all sequenced Ftc555-1 progeny isolates. Since the Idnurm laboratory has reported that the loss of *MLH1* results in elevated mutation rates [63], Ftc555-1 is likely to be a hypermutator strain. Increased mutation rates will drive phenotypic variations and some of those may be adaptive for survival in stressful environments, leading to rapid microevolution. On the other hand, the sequencing revealed that one gene (CNAG_03013; *OPT1*) was impacted by non-synonymous SNP changes and single nucleotide deletion in all three strain backgrounds. *OPT1* has been identified by Madhani group as an oligopeptide transporter required for transporting Qsp1, a quorum sensing peptide, into the receiving cells [53]. Deletion of *OPT1* exhibits similar phenotypes to our isolates, including increased capsule size and reduced melanin production, suggesting that this mutation may cause some of the phenotypic changes in our isolates. By reviewing the published sequences of 387 clinical and environments strains [64], we found that 6 of 287 clinical isolates contains high impact, potential loss of function mutations in *OPT1*, but no *OPT1* mutations in those 100 environmental isolates. The Fraser laboratory also reported that one of the clinical isolates in their study contains an inversion in chromosome 3 and affect two genes while one of them are *OPT1* [65]. The relatively high frequency of mutations in *OPT1* among clinical isolates suggests that this gene may be under particular selection during human infection. Another interesting gene mutation found in Ftc555-1 isolates was in the gene *PKR1*. This was a high impact mutation in F3, F4, F5 and F6, which exhibited phenotypes of titan cells and enlarged capsules inside macrophages and in macrophage medium. Pkr1 is known to be a negative regulator of titan cells and capsule enlargements in laboratory strains and clinical isolates [58,66]. A *pkr1* deletion mutant exhibit both enlarged capsule and titan cell production. It is also hypervirulent in a murine infection model [58].

The relatively low number of SNPs raises the question on how some of these strains change rapidly in response to amoeba predation that result in such broad and rapid phenotypic changes. Therefore, we also investigated the impact of whole-chromosome aneuploidy on isolates. An extra copy of chromosome 8 has been found in three isolates (H13, H16 and H17) which were isolated from the same pseudohyphal survival colony. Aneuploidy is caused by abnormal chromosomal segregation and can happen within even a single mitotic division, so this type of mutant can occur rapidly. This drastic DNA structural change often results in decreased fitness [67]. However, when fungi are exposed to stress, such as antifungal drugs, specific chromosomal aneuploidies can be advantageous through selection for increased gene expression of a subset of genes [55,68–72]. In *C. albicans* and *C. neoformans*, extra copies of specific chromosome containing drug resistance genes have been frequently found in antifungal drug resistance strains [55,70,71]. Likewise, *C. neoformans* could gain an extra chromosome as a solution for adaptation when the fungi encounter threats from amoebae. For instance, chromosome 8 contains one gene (*ZNF2*) which encodes a zinc-finger transcription factor that drives hyphal growth upon overexpression [73]. Chromosome 8 also contains another gene (*CBK1*) that is responsible for pseudohyphal formation [52,74]. *CBK1* encodes serine/threonine protein kinase which is one of the components of RAM pathway. Mutants in the RAM pathway have pseudohyphal phenotype, but we are not aware of any reports showing the effect of the overexpression of *CBK1* on pseudohyphae morphology. Since filamentous morphologies are important for resistance to phagocytosis by amoebae, it is possible that duplication of chromosome 8 could increase the cryptococcal fitness rapidly after exposure to amoebae. Indeed, when we re-introduced those aneuploid strains to amoebae, they could switch to filamentous forms quicker than their ancestor and efficiently resisted killing by amoebae. When we eliminated the chromosomal duplication, the phenotypes were restored back to wildtype level, supporting that there is strong link between duplication of chromosome 8 and amoebae resistance and other changes on virulence phenotypes such as capsule size and urease activity. In addition, there is no point mutations or structural changes such as amplification or deletion of chromosomal segments in these isolates. Therefore, aneuploidy may be the major source of the phenotypic change in that particular group of isolates.

Pseudohyphae are chains of elongated yeast cells that are unable to undergo cytokinesis completely, leading to multinuclei. Multinucleated cells showed a high level of chromosome instability, resulting in polyploidy and aneuploidy in eukaryotic cells [56]. Previous study of live-cell imaging on *Candida albicans* showed that hyphal cells occasionally generated multinucleated yeast cells [75], with polyploidy and/or aneuploidy, but there are very limited studies on whether pseudohyphal or hyphal formation may directly affect the ploidy variation. In this study, nuclear division, detected with GFP-H2B, was observed in cryptococcal pseudohyphae isolated from amoebae culture. The time-lapse imaging detected a nuclear fusion event, suggesting the cell experienced atypical nuclear division and potentially may undergo polyploidization which frequently generates their offspring with amplification of chromosomal segments or whole-chromosome aneuploidy. This result implies that interaction with amoebae not only contributes to the selection and maintenance of traits in *C. neoformans*, but also may drive heritable variation through pseudohyphae formation.

The ‘amoeboid predator-fungal animal virulence hypothesis’ formulates the notion that the capacity for virulence in soil fungi with no need for an animal host arose accidently from the traits for survival against ameboid predators that accidently also functioned as virulence factors for animal infection [12]. Consistent with this notion there is a remarkable concordance between fungal phenotypes that promote survival against amoeba and in animal hosts [14,23] and passage in amoeba is associated with increased virulence for several fungal species [24,39,76]. Analysis of virulence for the amoeba-selected strains described in our study in wax moths revealed no major changes in virulence from the parental strains. It is possible that this host does not discriminate between passaged and non-passaged *C. neoformans* cells or that none of the isolates tested gained or lost traits associated with virulence in that particular host. It is also possible that these strains already had the maximum pathogenic potential [77] for these animal hosts, which could not be further increased by amoeba interactions. However, we did observe that some amoeba-passaged strains were significantly more cytotoxic for macrophages *in vitro*. This result is consistent to the finding that which those strains also had great resistance to amoebae killing. The mechanism behind that is still unclear. However, those particular amoeba-passage strains can form larger cell size and capsule in both amoebae and macrophage culture and that may help them to escape from and cause damages to the host cells. These results fit the theory that amoebae are the training grounds for macrophage resistance of pathogens since the hostile environments in amoebae and macrophage are similar. Among these strains, the virulence of isolates F5 and F6 were further tested in murine infection model. These particular strains were picked because they acquired a mutation in *PKR1*, and deletion of *PKR1* has been shown to increase virulence [58]. However, neither F5 and F6 exhibited hypervirulence phenotype during murine infection, and instead were cleared faster than their parental isolate. It is noteworthy that the nonsense mutation found in F5 and F6 located in codon 407 which is only 75 codons prior to the original stop codon of *PKR1*. It is possible that the mutation results in altered function rather than loss of function and this is not sufficient to reproduce the hypovirulence phenotype caused by full *PKR1* knockout. Microbial virulence is a complex property that is expressed only in a susceptible host and host damage can come from the microbe or the immune response. Both F5 and F6 were able to establish themselves in the lung but triggered a more effective immune responses that cleared them. This finding implies the occurrence of other amoeba-selected changes that affect the immune response including overriding of the hypervirulence phenotype caused from the mutation of *PKR1* by compensation from other mutations or changes.

The amoeba-passaged *C. neoformans* selected in our study differ from those reported in prior studies [24,39,76] in that they did not increase in virulence. Instead, we observed reductions in murine virulence for two of the isolates studied despite increased capacity to damage macrophages from their long interaction with amoeba. Given the pleiotropic changes observed in our isolate set it is possible that we did not sample sufficient numbers to observed more virulent strains. Our study differs from prior amoeba-*C. neoformans* studies [39] in that it involved prolonged selection on a semi-solid agar surface in conditions that favored the protozoal cells by the presence of cations. In these conditions, amoeba dominance is manifested by a zone of fungal growth clearance where only occasional *C. neoformans* colonies emerged after several weeks. These colonies presumably emerged from resistant cells that survived the initial amoeba onslaught and gave rise to the variant strains that were analyzed in this study. We posit that these amoeba-resistant cells were very rare in the parent *C. neoformans* population and had emerged from the mechanisms discussed above, namely mutation and aneuploidy, which by chance conferred upon those cells amoeba resistance. Alternatively, those colony ancestor cells represent rare cells that were able to sense the amoeba danger and turn on diversity generating mechanisms that occasionally produced amoeba-resistant strains. In this regard, *C. neoformans* can sense amoeba and respond by increasing the size of its capsule by sensing protozoal phospholipids [40] but this process takes time and fungal cell survival probably depends on the race between adaptation and predation. The selection versus adaptation explanations for the origin of these are not mutually exclusive and both could have been operational in these experiments. These survivor cells then grew into a colony under constant amoeba selection where they gave rise to progeny cells where these phenotypic diversity generating mechanisms were maintained and amplified thus accounting for the phenotypic diversity observed in this study.

In summary, amoebae predation places great selective pressure in *C. neoformans* resulting in the rapid emergence of new phenotypes. The mechanism for these changes includes mutations and aneuploidy, which combine to create great phenotypic diversity. The effect of the phenotype diversification on the fitness of the fungi vary within the same or different hosts, which could promote fungal survival by a bet-hedging strategy that spreads the risk in situations where the environmental threat is unpredictable. Given that human infection also results in rapid fungal microevolution in this host, it is likely that similar mechanisms occur in vivo when this fungus comes under attack by immune cells. Indeed, several studies have shown microevolution of *Cryptococcus* during mammalian infection [65,78–80]. A bet hedging strategy that generates a prodigious number of phenotypes would increase survival in the face of unknown threats and could represent a general mechanism for survival in soils. Interference with the mechanism responsible for generating this plasticity could in turn result in new antimicrobial strategies that would reduce the emergence of diversity and thus simplify the problem for the immune response. Hence, it is interesting to hypothesize that amoeba predation in *C. neoformans* pushes a trigger that sets forth a series of events that generate diversity and similar mechanisms exist in other soil fungi that must routinely confront similar stresses.

## Method and material

### Ethics statement

All animal procedures were performed with prior approval from Johns Hopkins University (JHU) Animal Care and Use Committee (IACUC), under approved protocol numbers MO18H152. Mice were handled and euthanized with CO_2_ in an appropriate chamber followed by thoracotomy as a secondary means of death in accordance with guidelines on Euthanasia of the American Veterinary Medical Association. JHU is accredited by AAALAC International, in compliance with Animal Welfare Act regulations and Public Health Service (PHS) Policy, and has a PHS Approved Animal Welfare Assurance with the NIH Office of Laboratory Animal Welfare. JHU Animal Welfare Assurance Number is D16-00173 (A3272-01). JHU utilizes the United States Government laws and policies for the utilization and care of vertebrate animals used in testing, research and training guidelines for appropriate animal use in a research and teaching setting.

### Cell culture

*Acanthamoeba castellanii* strain 30234 was obtained from the American Type Culture Collection (ATCC). Cultures were clinical isolate of maintained in PYG broth (ATCC medium 712) at 25°C according to instructions from ATCC. *C. neoformans var. grubii* serotype A strain H99 and two environmental isolates A1-35-8 and Ftc555-1 were used for the interaction with amoebae, and these strains were originally obtained from John Perfect (Durham, NC). Both A1-35-8 and Ftc555-1 are environmental strains. A1-35-8 with genotype of VN1 molecular type is isolated from pigeon guano in US while Ftc555-1 is isolated from a mopane tree in Botseana and portrayed VNB molecular type. Both strains were avirulent in mouse model. Histone 2B-GFP tagged (C1746) H99 strain which was used for visualization of nuclear division of pseudohyphae was obtained from Kyung Kwon-Chung (Bethesda, MD) [81]. Cryptococcal cells were cultivated in Sabouraud dextrose broth with shaking (120 rpm) at 30°C overnight (16 h) prior to use in all experiments.

Bone-marrow derived macrophages (BMDM) were isolated from the marrow of hind leg bones of 5-to 8-wk-old C57BL-6 female mice (Jackson Laboratories, Bar Harbor, ME). For differentiation, cells were seeded in 100 mm TC-treated cell culture dishes (Corning, Corning, NY) in Dulbecco’s Modified Eagle medium (DMEM; Corning) with 20 % L-929 cell-conditioned medium, 10 % FBS (Atlanta Biologicals, Flowery Branch, GA), 2mM Glutamax (Gibco, Gaithersburg MD), 1 % nonessential amino acid (Cellgro, Manassas, VA), 1 % HEPES buffer (Corning), 1 % penicillin-streptomycin (Corning) and 0.1 % 2-mercaptoethanol (Gibco) for 6-7 days at 37 °C with 9.5 % CO_2_. Fresh media in 3 ml were supplemented on day 3 and the medium were replaced on day 6. Differentiated BMDM were used for experiments within 5 days after completed differentiation.

### Assay of *A. castellanii* and *C. neoformans* interaction

Two hundred *C. neoformans* yeast cells were spread on Sabouraud agar, and incubated at 30 °C overnight. *A. castellanii* in total 5 × 10^3^ cells were dropped randomly at several locations on the agar plate containing *C. neoformans*. Plates were sealed with parafilm and incubated at 25 °C for 3-4 months until survival colonies of *C. neoformans* emerged.

To isolate individual cell (hyphae or pseudohyphae in this case) out from the colony (Fig 1D), survival colonies were randomly picked from the plate to a 3 cm culture dish with PBS using pipette tips. Individual cells were picked under a light microscopy using pipette and transferred into a fresh Sabouraud agar. The plates were incubated at 30 °C. After 24 h incubation, the morphologies of microcolony were visualized using a Zeiss Axiovert 200M inverted microscope with a 10× phase objective. After 72 h incubation, colony morphologies were examined using Olympus SZX9 microscope with 1x objective and 32x zoom range. Morphologies of cells from colonies were visualized using Olympus AX70 microscope with 20x objective using the QCapture Suite V2.46 software (QImaging, Surrey, Canada).

### Amoebae killing assay

*C. neoformans* in 5 × 10^6^ cells were spread as a cross onto Sabouraud agar, and incubated at 30 °C for overnight. *A. castellanii* (10^4^) cells were dropped at the center of the *C. neoformans* cross. The plates were sealed in parafilm, and incubated at 25 °C. The distance from center to the edge of the clear predator zones in four directions was measured after 1-3 weeks incubation. The data were represented as the average of the distances of clear zone from four direction.

*C. neoformans* cells were also taken from the edge of the clear zone and at the end of the cross after 1-week incubation, and visualized using Olympus AX70 microscope with 20x objective. For samples of Ftc555-1 strains, the cells were counterstained with India ink.

### Capsule and cell size

*C. neoformans* cells were incubated in minimal medium (15 mM dextrose, 10 mM MgSO_4_, 29.4 mM KH_2_PO_4_, 13 mM glycine, 3 μM thiamine-HCl) at 30 °C for 72 h. In addition, Ftc555-1 and its isolates were incubated in medium for BMDM at 37 °C for 24 h. BMDM (1.5 × 10^6^ cells) were also infected with Ftc555-1 and its isolates (1.5 × 10^6^ cells) in 6-well plates. After 24 h infection, the culture supernatant was collected and the plates were washed once to collect the extracellular *C. neoformans*. The intracellular *C. neoformans* was collected by lysing the host cell with sterile water. The cells were stained with 0.1% Uvitex 2B (Polysciences, Warrington, PA) for 10 min and washed two times with PBS. The capsule was visualized by India ink negative staining by mixing cell samples with equal volume of India ink on glass slides and spreading the smear evenly with coverslips. The images with a minimum 100 randomly chosen cells was taken by using Olympus AX70 microscopy with 40x objective at bright-field and DAPI channel. The areas of cell body and whole cell (cell body plus capsule) were measured using image J software. The capsule thickness was calculated by subtracting the diameter of whole cell from that of cell body. The cell size was presented as the diameter of cell body without capsule. Three biological independent experiments were performed for each sample.

### lactate dehydrogenase (LDH) release assay

BMDM cells (5 × 10^4^ cells/well) were seeded in 96-well plates with BMDM for overnight. To initiate the phagocytosis, *C. neoformans* with 5 × 10^5^ cells in the presence of 10 µg/ml 18B7 mAb were added in each well of BMDM culture. The culture plates were centrifuged at 1200 rpm for 1 min to settle yeast cells on the monolayer of macrophage culture. After 48 h infection, LDH release were assessed using CytoTox-ONE Homogeneous Membrane Integrity Assay kit (Promega, Madison, WI) according to the manufacturers’ instructions.

### Urease activity

*C. neoformans* in 10^8^ cells were incubated in 1 ml of rapid urea broth (RUH) developed by Roberts [82] and adapted by Kwon-Chung [83] at 30 °C. After 1-4 h incubation, cells were collected by centrifugation and 100 µl of supernatant were transferred to 96-well plate. The absorbance of the supernatant was measured at 570 nm using EMax Plus microplate reader (Molecular Devices, San Jose, CA). The assay was performed in Triplicate for each time interval.

### Melanin quantification

*C. neoformans* in 10^4^, 10^5^, 10^6^ and 10^7^ cells were spotted on agar of minimal medium supplemented with 1 mM L-DOPA (Sigma Aldrich, St Louis, MO). The plates were incubated at 30 °C without light. Photos were taken after 1-3-day incubation on a white light illuminator. The photos of samples were always taken together with their ancestors under the same condition in order to avoid different exposure time or light adjusted by the camera. The obtained photos were then converted to greyscale using image J software. The regions of the colonies were selected and the pixels of each selected region were quantified in grayscale. The relative grayscale of the colonies from samples were normalized by the grayscale of the colonies of ancestors. The representation data shown in this paper are at the cell number of 10^6^ cells and at the time point of 24 h. Three biological independent experiments were performed for each sample.

### Macrophage killing assay

BMDM cells (5 × 10^4^ cells/well) were infected with *C. neoformans* (5 × 10^4^ cells) in the presence of 10 µg/ml 18B7 mAb. The culture plates were centrifuged at 1200 rpm for 1 min to settle yeast cells on the monolayer of macrophage culture. After 24 h infection, phagocytized cryptococcal cells were released by lysing the macrophages with sterilized water. The lysates were serially diluted, plated onto Sabouraud agar and incubated at 30 °C for 48 h for colony form unit (CFU) determination. This experiment was performed in triplicates for each strain.

### Virulence assay in *Galleria mellonella*

*G. mellonella* larvae were purchased from Vanderhorst Wholesale (Saint Mary’s, OH). Larvae were picked based on weight (175 – 225 mg) and appearance (creamy white in color). Larvae were starved overnight at room temperature. Next day, overnight cultures of *C. neoformans* that grew in Sabouraud broth were washed three times with PBS and diluted to 1×10^5^ cells/ml. Cells in 10 µl were injected into the larva via the second last left proleg paw with 31G needles. Infected larvae were incubated at 30 °C and the number of death larvae were scored daily until all the larvae infected with *C. neoformans* ancestral strains in this study were dead. Control groups of larvae were inoculated with 10 μL of sterile PBS. Experiments were repeated at least two times with experimental groups of 15 larvae at a time.

### Whole genome sequencing and variant identification

Genomic DNA was prepared using cetyltrimethylammonium bromide (CTAB) phenol-chloroform extraction as described previously [84]. Genomic DNA was further purified using a PowerClean DNA cleanup kit (QIAGEN, Hilden, Germany). Libraries were constructed using the Illumina DNA Flex Library kit and were sequenced on an Illumina HiSeq2500 to generate paired 150 base reads. An average of 145X sequence depth (range 69-176X) was generated for each sample. All sequence for this project is available in NCBI under BioProject PRJNA640358.

Reads were aligned to the *C. neoformans* H99 assembly [85] using BWA mem v0.7.12 [86]. Variants were identified using GATK v3.7 [87]; HaplotypeCaller was invoked in GVCF mode with ploidy = 1, and genotypeGVCFs was used to predict variants in each strain. The workflow used to execute these steps on Terra (terra.bio) is available on Github (https://github.com/broadinstitute/fungal-wdl/tree/master/workflows/fungal_variant_calling_gatk3.wdl). Sites were filtered with variantFiltration using QD < 2.0, FS > 60.0, and MQ < 40.0. Genotypes were filtered if the minimum genotype quality < 50, percent alternate allele < 0.8, or depth < 10 (https://github.com/broadinstitute/broad-fungalgroup/blob/master/scripts/SNPs/filterGatkGenotypes.py). Genomic variants were annotated and the functional effect was predicted using SnpEff v4.1g [88].

### Cryptococcal cell karyotyping

Cell karyotypes were analyzed by quantitative PCR. qPCR primers used in this study have been published in Gerstein et al. 2015. qPCR reactions were performed in a StepOnePlus Real-Time PCR System (Applied Biosciences, Beverly Hills, CA) using 20 µl reaction volumes. All reactions were set up in technical triplicate. Each reaction mixture contained PowerUp SYBR Green Master Mix (Applied Bioscience), 300 nM each primer, 10-ng genomic DNA from CTAB extraction, and distilled water (dH2O). Cycling conditions were 95°C for 5 min followed by 40 cycles of 95°C for 15 s and 55°C for 1 min. Melt curve analysis was performed in 0.5°C increments from 55 to 95°C for 5 s for each step to verify that no primer dimers or product from misannealed primers had been amplified. Threshold cycle (CT) values were obtained using StepOnePlus software version 2.3 (Applied Bioscience) where the threshold was adjusted to be within the geometric (exponential) phase of the amplification curve. Chromosome copy numbers were determined using a modified version of the classical CT method as described by [69].

### Visualization of nuclear division in pseudohyphae

Histone 2B-GFP tagged H99 (C1746) was interacted with *A. castellanii* on Sabouraud agar as described above until survival colonies with pseudohyphae emerged. The colonies were transferred on the well of 18B7 Ab coated coverslip bottom MatTek petri dishes with 14mm microwell (MatTek Brand Corporation, Ashland, MA) in minimal medium. After 30 min incubation to allow for settling down the cells, 2 ml of minimal medium were added. Images were taken every 10 min for 24 h using of a Zeiss Axiovert 200M inverted microscope with a 10x phase objective and GFP channel in an enclosed chamber under conditions of 30 °C.

### Measurement of global histone H3 acetylation

A culture of *C. neoformans* in 2 ml was grown in Sabouraud broth for 24 h. Protein samples were extracted by vortexing for 4 h with 0.5 mm glass beads and yeastbuster extraction buffer (Merck, Darmstadt, Germany) at 4 °C. Supernatant were collected and 100% trichloroacetic acid was added at a 1:4 ratio. The mixtures were incubated on ice for 30 min, and pellets were then collected by centrifugation at 13,000 g for 10 min at 4°C. Pellets were washed twice with 1 ml acetone and dissolved in 20 µl water. The protein concentrations were measured by using Micro BCA(tm) Protein Assay Kit (Themofisher, Waltham, Ma). The protein samples (3.5 µg) were used to detect the global histone H3 acetylation levels by using the EpiQuik global histone H3 acetylation assay kit (EpigenTek, Farmingdale, NY) according to manufacturer’s instructions.

### Stress sensitivity test

The overnight cultures were diluted in Sabouraud broth to an OD_600_ of 2 and further diluted 10-, 10^2^-, 10^3^-, 10^4^-, 10^5^-fold. The dilutions (5 μl) were spotted onto Sabouraud agar plates supplemented with 16 µg/ml fluconazole and incubated for 48 h at 30 °C. Plates without fluconazole were also incubated for 48 h either 30 or 37 °C.

### Growth curve

*C. neoformans* strains Ftc555-1, F5 and F6 were grown in Sabouraud media at 30°C with orbital shaker (120 rpm) for 7 days with data measurements each 24 hours. The assay was performed in a 96-well plate and some serial dilutions were done, with a cell concentration range between 1.0 × 10^7^ to 5.0 × 10^3^/well. Each condition was done in triplicate. The growth was measured by optical density at 600 nm.

### Murine Infection

Six-week-old female A/J mice were infected intranasally with 20 µl of 1.0 × 10^7^ yeast cells of each *C. neoformans* strain. Three groups of mice (n = 8 animals per group) were infected and deaths were scored daily for 60 days. No death was observed during this time, so we decided to euthanize the animal for fungal burden assessment and cytokines levels determination. A second experimental infection was performed with some modifications. Six-week-old female A/J mice were infected intranasally with 20 µl of 1.0 × 10^7^ yeast cells of each *C. neoformans* strain (n = 5 animal per group) and then euthanized after 5 days. Specific organs were removed for fungal burden and cytokines level evaluation.

### Fungal burden assessment

The fungal burden was measured by counting CFU (colony-forming units). After animals euthanasia, the lungs were removed, weighed and homogenized in 1 ml of PBS. After serial dilutions, homogenates were inoculated onto Sabouraud agar plates with 10 U/ml of streptomycin/penicillin. The plates were incubated at room temperature, and the colonies were counted after 48-72 h.

### Determination of cytokine levels in the organs

Spleen and lungs of each mouse were macerated with protease inhibitor (complete, EDTA-free, Roche Life Science, Indiana, United States) and centrifuged; supernatants of these samples were used for cytokine detection by a sandwich-ELISA by using commercial kits (BD OptEIA(tm), BD Franklin Lakes, Nova Jersey, US) for the following cytokines: IL-2 (#555148), IL-4 (#555232), IL-10 (#555252), IFN-γ (#551866) and TNF-α (#555268).The protocol was followed according to the manufacturer’s recommendations. The reading was performed in a plate spectrophotometer at 450 nm and 570 nm.

## Acknowledgements

We thank the Broad Institute Microbial Omics Core for generating the DNA libraries and the Genomics Platform for the sequence for this study.

## Supporting information

S1 Table. High impact indels found in passaged Ftc555-1 isolates

S1 Fig. Relative chromosome copy number of isolate H14 was obtained by qPCR.

S2 Fig. Amoebae killing assay on *C. neoformans* deletion mutants. Mutants showed comparable predator zone with their parental strains.

S3 Fig. Global histone H3 acetylation levels in parents and isolates. Data are means of two independent experiments with standard deviations.

S4 Fig. The growth curves of Ftc555-1, F5 and F6 strains with high (1.0×10^7^) and low (5.0×10^3^) inoculum concentration in Sabouraud medium for seven days.

S5 Fig. Cytokine production in spleen of mice after infection with Ftc555-1 as well as its isolates F5 and F6. After 60 and 5 days of infection, mice were sacrificed and the level of cytokines (A, E) IL4 (B, F) IL10 (D) IFN-γ (D, G) TNF-α in the lung were measured by ELISA. All data represent the mean of eight mice per group and errors bars are SD. For determination of cytokine levels, one-way ANOVA with Kruskal-Wallis nonparametric test was used and followed by Bonferroni’s multiple-comparison test. The t-test was used to compare the number of colony forming units (CFU) for different groups.

S6 Fig. Virulence of parents and variant isolates in the *G. mellonella* larvae infection model. The Kaplan-Meier plots shows the survival of *G. mellonella* after injection of cryptococcal cells (10^3^ cells/larva).

S7 Fig. Summary of phenotypic changes occurred in amoeba-passaged isolates S1 Movie. Time-lapse imaging showing nuclear division of pseudohyphae.

